# Droplet printing reveals the importance of micron-scale structure for bacterial ecology

**DOI:** 10.1101/2020.10.20.346577

**Authors:** Ravinash Krishna Kumar, Thomas A. Meiller-Legrand, Alessandro Alcinesio, Diego Gonzalez, Despoina A. I. Mavridou, Oliver J. Meacock, William P. J. Smith, Linna Zhou, Wook Kim, Gökçe Su Pulcu, Hagan Bayley, Kevin R. Foster

## Abstract

Bacteria often live in diverse communities where the spatial arrangement of strains and species is considered critical for their ecology, including whether strains can coexist, which are ecologically dominant, and how productive they are as a community^1,2^. However, a test of the importance of spatial structure requires manipulation at the fine scales at which this structure naturally occurs^3–8^. Here we develop a droplet-based printing method to arrange different bacterial genotypes across a sub-millimetre array. We use this to test the importance of fine-scale spatial structure by printing strains of the gut bacterium *Escherichia coli* that naturally compete with one another using protein toxins^9,10^. This reveals that the spatial arrangement of bacterial genotypes is important for ecological outcomes. Toxin-producing strains largely eliminate susceptible non-producers when genotypes are well-mixed. However, printing strains side-by-side creates an ecological refuge such that susceptible strains can coexist with toxin producers, even to the extent that a susceptible strain outnumbers the toxin producer. Head-to-head competitions between toxin producers also reveals strong effects, where spatial structure can make the difference between one strain winning and mutual destruction. Finally, we print different potential barriers between two competing strains to understand why space is so important. This reveals the importance of processes that limit the free diffusion of molecules. Specifically, we show that cells closest to a toxin producer bind to and capture toxin molecules, which creates a refuge for their clonemates. Our work provides a new method to generate customised bacterial communities with defined spatial distributions, and reveals that micron-scale changes in these distributions can drive major shifts in their ecology.

---

Bacteria commonly exert strong effects on the growth and survival of surrounding cells, via mechanisms including nutrient competition, quorum sensing, and toxin production^11–14^. Consequently, the position of different strains and species in space is thought to be critical for the ecology of bacterial communities, and indeed how they affect us^1,5–8^. In particular, the way in which different strains of bacteria are arranged when a community first develops is expected to be a key predictor of which strains dominate and, more generally, whether the community thrives or perishes^15–24^. When strains and species grow apart in distinct patches, for example, this is expected to limit the impacts of competitive mechanisms, thereby promoting both coexistence and productivity. The importance of spatial structure is broadly supported by comparisons of liquid with solid-surface (agar) culture, and agar-based manipulations on the millimetre to centimetre scale (Fig. 1b)^25–30^. However, natural spatial structure often occurs at much finer scales, with data from the mammalian microbiome suggesting that the micron to millimetre scale is particularly important^3,6,31^. For example, while there appears to be significant genotypic mixing in the gut microbiome, the dental and tongue microbiome is more structured, with patches of bacteria dominated by one genotype reaching hundreds of microns in scale^5,8^. Moreover, there is evidence that solute gradients within bacterial communities can limit the distance at which cells affect each other to micron scales^32–34^, raising the possibility that differences in the arrangement of genotypes at these scales will be important.

**Figure 1.**
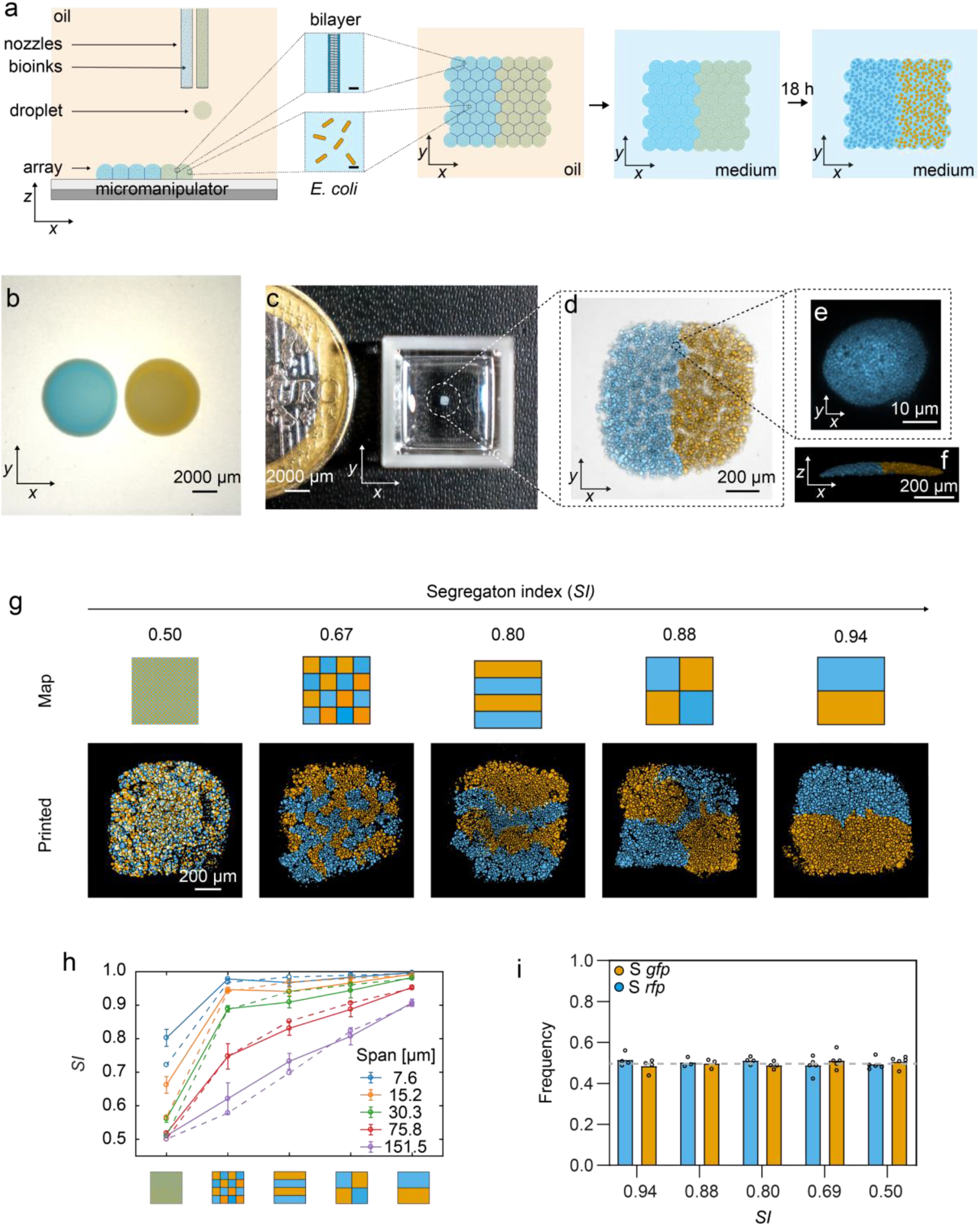
Droplet-printing can manipulate micron-sized spatial structure when forming bacterial communities. **a**, A schematic of the fabrication of printed bacterial communities by droplet-printing (depicted in the *x, z*, and *x, y* dimensions), transfer to LB medium, and growth. **b**, A composite (brightfield and fluorescence) stereomicroscope image of two side-by-side spotted colonies of susceptible (S) *gfp* and S *rfp* strains grown for 24 h at 37 °C from starting cell densities of 10^8^ cells mL^-1^. **c**, A photograph of a printed array (after 18 h of growth and transfer to M9 medium) with a 1-euro coin for size comparison. Images in **b** and **c** are taken at the same magnification. **d**, A higher magnification, composite (transmitted light and fluorescence) confocal microscope image of a 7 x 8 x 1 (*x, y, z* droplets) printed array containing S *gfp* and S *rfp* strains segregated to opposite sides of the structure. **e**, A confocal fluorescence microscopy image of an S *rfp* microcolony in the *x*, *y* planes. **f**, A 3D rendering of *z*-stacked confocal microscopy images of the same printed community as in **d. g**, Segmented fluorescence images of printed communities (7 x 8 x 1 (*x, y, z* droplets)) of S *gfp* and S *rfp* strains at graded degrees of genetic mixing accompanied by the corresponding printing map and the calculated segregation indices (*SI*) at a local neighbourhood of 100 μm (see Methods). The local neighbourhood is the spatial scale on which genetic mixing is measured. In **b** to **g**, S *gfp* and S *rfp* strains are false-coloured orange and blue, respectively. **h**, A plot showing the calculated *SI* of printed arrays (smooth lines) at different sizes of local neighbourhoods (‘spans’) (7.6–151.5 μm) compared to the *SI* of the printing maps (dashed lines). Data points are the mean of *n* = 9 printed arrays and error bars are the standard deviation. **i**, A bar chart showing the frequency of S *gfp* and S *rfp* strains in arrays printed at different *SI* values. Each data point is a biological replicate and the height of the bars are the mean frequencies of the strains. The grey line shows the initial starting frequency (0.5 for both genotypes). A Kruskal-Wallis test found a non-significant difference between the median frequencies of S *gfp* at different *SI* values (P = 0.2736; see Supplementary Table 1 for statistical tests). In **i**, the term ‘frequency’ (*f*) of a genotype *a* in a printed array containing genotypes *a* and *b* after 18 h of competition is defined as: *f = A_a_*/(*A_a_* + *A_b_*), where *A* is the total cross-sectional area that a genotype occupies in the printed array.

A test of the importance of spatial structure requires the ability to manipulate the distribution of different bacteria in communities at fine spatial scales. To enable this test, we developed a new droplet printing method that allows us to position microbes in specific patterns in sub-millimetre arrays (Fig. 1). The method works by piezo injection of tailored ‘bioinks’ comprising bacterial cells (*E. coli*, ≈ 10^8^ cells mL^-1^), molten agarose (1.5% w/v), and lysogeny broth (LB) medium into an oil/phospholipid solution at 37°C. The method generates 110 μm-diameter droplets (≈70 cells per droplet), which are deposited into 2D patterns by line-by-line printing (Fig. 1a). The droplets are initially surrounded by monolayers of phospholipid, which form bilayers on contact with one another, thereby creating a stabilised, support-free structure^35,36^ (Fig. 1a). The resulting array is then gelled, and the lipid bilayers removed by the addition of an oil (silicone oil AR20, which causes the lipid to precipitate) to create a final printed community of bacteria in agarose suspended in LB medium (Methods) (Fig. 1a, c–f). Within the printed community, bacteria grew from single-cell dispersions (with starting 2D centre-to-centre distances of 18 ± 2 μm) into 3D microcolonies (with median diameters of 15 (10–21) μm) over 18 h of growth (see Extended Fig. 1 and Supplementary Methods 1 for further characterisation of printed arrays). To validate the method, we printed two fluorescently labelled strains of *E. coli*. This demonstrated the ability to print specific patterns and, with them, to vary the amount of genetic mixing (defined by a segregation index (*SI*)^19^, see Methods and Extended Fig. 2) at the scale of the arrays (Fig. 1g). Importantly, the calculated *SI* of our printed arrays matched well with the theoretical *SI* of the respective printing maps (Fig. 1h), and the imposed patterns did not impact the frequencies of strains that differ only in the fluorophore that they carry (Fig. 1i) (Extended Fig. 2 and Supplementary Methods 1.2). With this method, therefore, we can systematically alter the spatial arrangement of bacteria strains across naturally-relevant scales to study how they interact and grow.

We next sought to test whether this spatial structure is important for the ecology of interacting bacteria. To do this, we studied a common ecological interaction in the mammalian gut: interference competition between strains of *E. coli* that produce protein toxins known as colicins^10,27,37^. While colicin production can be critical for survival, there is diversity in the types of colicins carried by *E. coli* strains, and many strains carry none at all — suggesting that colicin release is not always beneficial^38^ (Supplementary Methods 2.1). We began by performing two-way competitions between susceptible non-colicin producing strains (S) and a range of colicin-producing strains (P) in printed communities at different *SI* values (Fig. 2) (see Table 1 for strains used). If fine-scale structure is important, theory suggests that genetic mixing will tend to favour colicin producers, as it gives them more access to susceptible cells. Conversely, increasing spatial structure has the potential to isolate producer genotypes from susceptible genotypes, thereby protecting them (Fig. 2b, c)^39–41^. However, it was far from clear whether micron-scale spatial structure would have any effect because colicins are released from cells (they do not remain attached)^42^, will diffuse through agar across centimetre scales^29,42^, and are highly toxic, killing with single-hit kinetics^43,44^.

**Figure 2.**
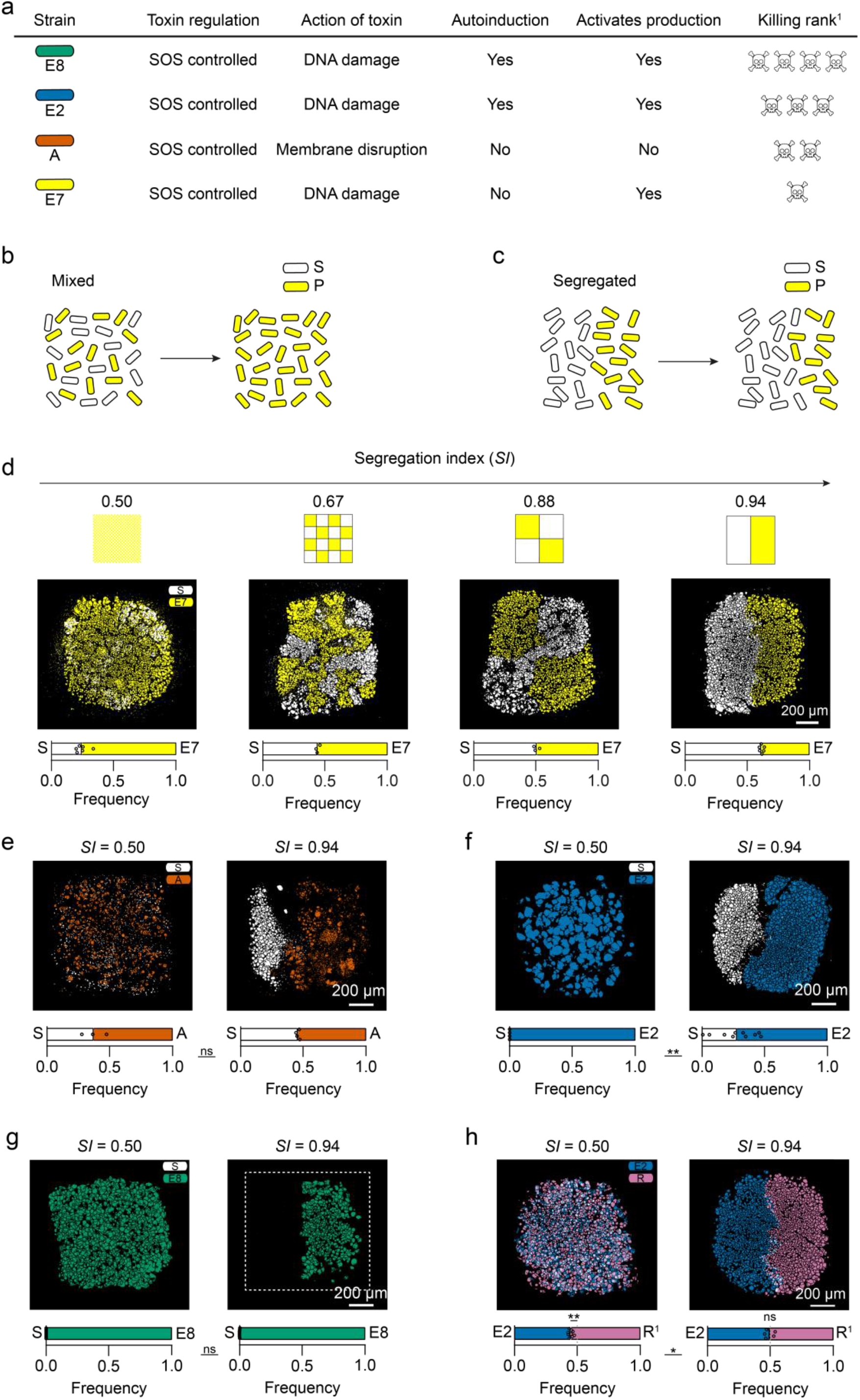
Micron-scale structure is a critical determinant of the outcome of interference competition between susceptible and colicin-producing strains. **a**, A diagram describing the characteristics of each colicin-producing strain used in this study. ^1^The ‘killing rank’ ranks the ability of each of the colicin-producing strain to kill colicin-susceptible strains, see [42 & 45] and data therein. **b–c**, A schematic hypothesis, of how susceptible and colicin-producer frequencies change over time, in a genetically-mixed spatial structure (*SI* = 0.50) (**b**), and in a genetically segregated spatial structure (*SI* = 0.94) (**c**), if susceptible growth rates outweigh the cost of toxin-production. **d**, Segmented fluorescence images of S cells (white) and E7 cells (yellow) in communities printed at an initial *SI* of 0.50 (*n* = 6), 0.67 (*n* = 3), 0.88 (*n* = 3), and 0.94 (*n* = 6) with accompanying printing maps. A Kruskal Wallis test showed a statistically significant relationship between the change in *S_Frequency_* with *SI* (Kruskal Wallis statistic: 15.63, *p* < 0.001, see Supplementary Table 2 for statistical test). **e–g**, Segmented fluorescence images of S cells (white) and A cells (orange) (*n* = 3 and *n* = 4 for *SI* = 0.50 and 0.94, respectively) (**e**), and E2 cells (blue) (*n* = 4 and *n* = 10 for *SI* = 0.50 and 0.94, respectively) (**f**), and E8 cells (bluish green) (*n* = 7 and *n* = 7 for *SI* = 0.50 and 0.94, respectively) (**g**), in communities printed at an initial *SI* = 0.50 and 0.94. **h**, Segmented fluorescence images of E2 cells (blue) and R^1^ cells (pink) in communities printed at an initial *SI* of 0.50 (*n* = 9) and 0.94 (*n* = 6). In **d–h**, each fluorescently segmented image was taken after 18 h of competition. Below each fluorescently segmented imaged is a stacked bar chart of the frequencies of strains in the array. Each data point represents a biological replicate; the mean frequencies of each strain are where the bars meet. In **e–h**, unpaired Mann Whitney tests were performed to assess the statistical significance of the changes in frequencies of competing strains at different *SI* values. For **e**, P = 0.6286; for **f**, P = 0.0020; for **g**, P = 0.4462; for **h**, P = 0.0336 (see Supplementary Tables 3 for statistical tests). In **h**, One sample t and Wilcoxon tests showed a statistically significant (P = 0.0039) and non-significant (P > 0.6250) difference between the median frequencies of E2 and R^1^ at *SI* = 0.50 and 0.94, respectively (see Supplementary Tables 3 for statistical tests). In **d–g**, the term ‘frequency’ (*f*) of a genotype *a* in a printed array containing genotypes *a* and *b* after 18 h of competition is defined as: *f* = *A_a_*/(*A_a_* + *A_b_*), where *A* is the total cross-sectional area that a genotype occupies in the printed array.

**Table 1.**
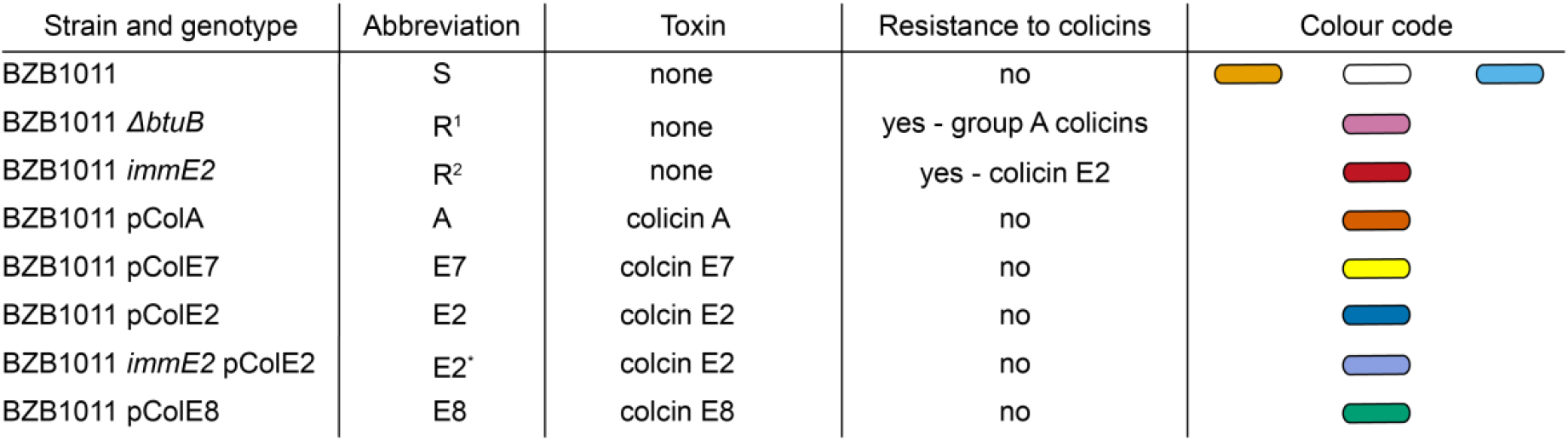
Strains used in this study. Each colicin-producing strain is immune to its cognate toxin. For BZB1011 (S), the orange and sky-blue colour codes are used to represent S cells expressing sfGFP and mRFP1, respectively, in Fig. 1 and 4. The white colour code represents S cells in Fig. 2 and 3.

| Strain and genotype | Abbreviation | Toxin | Resistance to colicins | Colour code |
| --- | --- | --- | --- | --- |
| BZB1011 | S | none | no |  |
| BZB1011 $\Delta btuB$ | R<sup>1</sup> | none | yes - group A colicins | |
| BZB1011 *immE2* | R<sup>2</sup> | none | yes - colicin E2 |  |
| BZB1011 pColA | A | colicin A | no |  |
| BZB1011 pColE7 | E7 | colicin E7 | no |  |
| BZB1011 pColE2 | E2 | colicin E2 | no |  |
| BZB1011 *immE2* pColE2 | E2<sup>*</sup> | colicin E2 | no |  |
| BZB1011 pColE8 | E8 | colicin E8 | no |  |

We found that altering spatial structure across the micron-scale had significant effects on the outcome of competition between the bacteria. For all but one of the colicin types (E8), the introduction of fine-scale spatial structure enabled a distinct patch of the susceptible strains to persist (Fig. 2d–h). The strength of this effect reflects the relative ability of each colicin strain to kill susceptible strains^42,45^ (Fig. 2a, Extended Fig. 3d-f). As might be expected, the more potent killers were less affected by spatial structure, with the colicin E8 producer killing off the susceptible cells irrespective of whether there is structure (Fig. 2g). E8 is known to be both a potent colicin and one that is produced at high levels due to autoinduction, whereby producing cells promote toxin production in clonemates (Extended Fig. 3e)^42,45^. For all the other strains, we found that spatial structure creates an ecological refuge^46^ for susceptible cells. Even though the refuge is only a few hundred microns across, it allowed susceptible cells to better survive competition with toxin-producing strains for two of the combinations. For E7, this effect was so strong that the susceptible strain was able to outgrow the colicin producer when printed side by side (Fig. 2d, S_*Frequency*_ = 0.65 ± 0.02 at *SI* = 0.94). Counts of colony-forming units confirmed that the imaged susceptible cells in the printed arrays were viable, despite the presence of the toxin producer (Extended Fig. 3i). Exploring the case of E7 in more detail revealed a graded response to spatial structure, where progressively decreasing spatial structure (decreasing *SI*) led to a progressive increase in the abundance of the E7 colicin producer (Fig. 2d). At high genotypic mixing, the outcome of the competition was reversed, where now the toxin producer outcompeted the susceptible strain (Fig. 2d, S_*Frequency*_ = 0.24 ± 0.05 at *SI* = 0.50).

We also found an effect of spatial structure when a colicin producer strain competes with a colicin-resistant strain (R^1^) rather than a susceptible strain (Fig. 2h). The colicin producer has a lower growth rate due to the costs of colicin production (Extended Fig 3g–h). However, this growth disadvantage only translates to an ecological disadvantage in the absence of spatial structure, i.e. when the two strains are well mixed (E2_*Frequency*_ = 0.46 ± 0.02 at *SI* = 0.50). When growing side-by-side in a structured community, the colicin producer reaches the same abundance as the resistant strain, despite its slower growth rate. We saw the same effect of spatial structure for another strain combination (S and R^1^) that lack any toxin mediated competition (Extended Fig. 4c). While our focus is interference competition (via toxins), these experiments revealed that exploitative competition (mediated by differences in growth rate) is also affected by the introduction of spatial structure in the printed arrays.

We see, therefore, that the arrangement of bacteria at the sub-millimetre scale is important for the outcome of both interference and exploitative competition. Moreover, as predicted by both ecological and evolutionary theory, our results suggest that the key effect of printing strains side by side – as opposed to well mixed – is to weaken interactions (Fig. 2b, c)^26,47,48^. We next explored the importance of positioning for competition between pairs of toxin-producing strains, which are each immune to their own toxin but able to kill the other (Fig. 3). At first, these appeared to follow the same pattern as the previous (producer vs susceptible) experiments. In one (E2 and E7), we saw the same refuge effect with spatial structure, where a strain that is eliminated under mixed conditions can persist when printed side by side with the more dominant strain (E2) (Fig. 3a). The dominance of E2 is expected because a higher proportion of E2 cells make colicins than E7 cells due to autoinduction^49,50^. In another case, one strain (A) eliminated the other (E2) irrespective of spatial structure (Fig. 3b). The potency of colicin A cells is expected because E2 is a DNase that activates the SOS response, which activates colicin A expression, but A does not active E2 expression because colicin A targets membrane integrity rather than DNA (Fig. 2a and Extended Fig. 3f)^29,51,52^. However, a third combination (E2 and E8) had a surprising outcome (Fig. 3c). Here, without spatial structure, the dominant strain E8 eliminated E2 as though E2 were simply a susceptible strain. By contrast, with spatial structure (*SI* = 0.94), both strains were almost completely eliminated.

**Figure 3.**
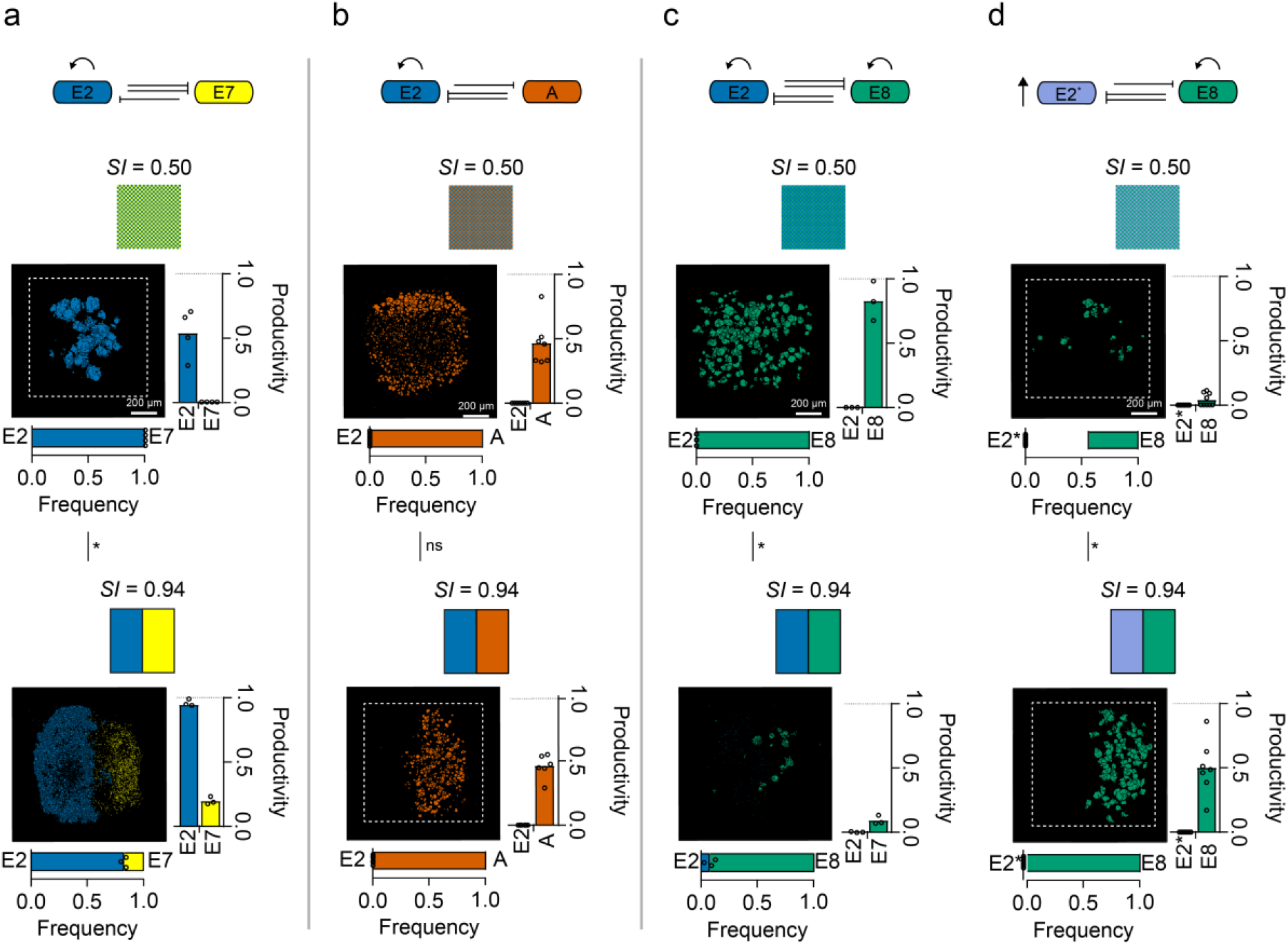
Micron-scale structure is a critical determinant of the outcome of interference competition between colicin-producing strains. **a–d**, Competition schematics followed by corresponding segmented fluorescence images of E2 cells (blue) and E7 cells (yellow) (*n* = 4 and *n* = 3 for *SI* = 50 and 0.94, respectively) (**a**), E2 cells and A cells (orange) (*n* = 7 and *n* = 6 for *SI* = 0.50 and 0.94, respectively) (**b**), E2 cells and E8 cells (bluish green) (*n* = 3 and *n* = 3 for *SI* = 0.50 and 0.94, respectively) (**c**), and E2* cells (bluish grey) and E8 cells (*n* = 9 and *n* = 7 for *SI* = 0.50 and 0.94, respectively) (**d**) in genetically-mixed (*SI* = 0.50) and fully segregated (*SI* = 0.94) printed communities after 18 h of competition. In the bacterial warfare schematics, the curly arrows represent autoinduction, the single inhibition arrows represent lower aggression, the double inhibition arrows represent higher aggression, and the vertical arrow represents a faster growth rate. Above each segmented fluorescence image is the corresponding printing map. Below each segmented fluorescence image is a stacked bar chart of the frequencies of strains in each printed array after 18 h of competition. Each data point is a biological replicate and the mean frequencies of each strain are denoted where the bars meet. The mean frequency of E8 in E2* vs E8 at *SI* = 0.50 (d) is shown as the length of the bar and the data points denote *E2*\*_*Frequency*_. To the right of each segmented fluorescence image is a bar chart of the productivities of strains in printed arrays. Each data point is a biological replicate and the mean productivities of each strain are denoted by the heights of the bars. In **a–d**, unpaired Mann-Whitney tests were performed to assess the statistical significance in the changes in frequencies of competing strains at different *SI* values. For **a,** P = 0.0286; for **b**, P > 0.9999; for **c**, P = 0.0286; and for **d**, P = 0.0337 (see Supplementary Tables 4 for statistical tests). In **a–d**, the term ‘productivity’ (*p*) of a genotype *α* in a printed array after 18 h of competition is defined as: *p* = *A_a_/A_S_*, where *A_S_* is the total crosssectional area that a susceptible genotype occupies after 18 h of competition (at the same starting density and spatial pattern as genotype *a*), but without interference competition.

We next sought to understand how spatial structure could enhance competition, and moreover do it to the extent that both populations collapse. We hypothesised that the answer lay in the fact that both of these strains exhibit autoinduction of colicin production. Similar to the effects of quorum sensing, this density-dependent regulation occurs because the import of a clonemate’s DNA-damaging colicin increases a cell’s probability of colicin release (Extended Fig. 3e). Autoinduction can have strong effects on competition between colicin strains^42^. In our experiments, we hypothesised that spatial structure provided E2 cells enough of a refuge to reach a sufficient cell density to activate collective colicin production through autoinduction, thereby allowing them to inhibit E8 cells more strongly. To explore this, we studied competitions between E8 and an E2 strain engineered to lack autoinduction in colicin production (E2*, see Methods) (Fig 3d). As predicted, this greatly reduced the impact of E2 on E8 in the presence of spatial structure. Moreover, the effects of spatial structure were reversed, with mutual destruction of the two competitors now occurring under mixed conditions without spatial structure. This ability of the engineered E2* strain to eliminate E8 under mixed conditions is likely to stem from the relatively high growth rate of E2* (Extended Fig. 3g, h), which allows it to rapidly establish itself in mixed arrays. In sum, the impact of spatial structure depends on whether E2 does, or does not, perform autoinduction. Moreover, in both cases, we see a strong effect where the printed pattern determines whether one strain thrives or both are largely eliminated.

The impacts of autoinduction demonstrate that fine-scale structure can have surprising and complex effects on bacterial interactions. Nevertheless, one effect that is commonly seen is that spatial structure creates a refuge, with subordinate strains better able to persist with spatial structure than without it. We sought, therefore, to understand the mechanistic basis of this refuge effect, using prints of three different types of bioinks to explore the interface between a colicin E2 producer and a susceptible strain (Fig. 4a). Printing cell-free agarose between an E2 strain and a susceptible strain removed the refuge effect, suggesting that the distribution of colicins was not limited by diffusion across the printed array (Fig. 4e). This result implied that the susceptible cells closest to the E2 producer were somehow interrupting the diffusion of colicins across the array. We hypothesised that this could occur if susceptible cells bind and thereby ‘mop up’ colicin molecules as they pass. To explore this, we engineered a colicin-resistant strain that lacks the BtuB receptor for colicin E2 (Table 1, Fig 4b, Extended Fig. 4a) (see Supplementary Methods 2.2). As predicted by our model, the refuge effect was eliminated even though there were cells growing next to the colicin producer (Fig. 4f). However, the BtuB mutation also makes the cells resistant to colicin (Extended Fig. 4b), such that the lack of protection could be linked in some way to the lack of cell death from E2 activity. To exclude this possibility, we engineered a strain that carries BtuB, but which is resistant to colicins via cytoplasmic expression of the E2 immunity protein (Table 1, Fig. 4b, and Extended Fig. 4a). This strain was not affected by the colicin of the E2 producer (Extended Fig. 4b) but did provide protection (Fig. 4g). We conclude, therefore, that binding of diffusing colicins by intoxicated cells is critical to the refuge effect.

**Figure 4.**
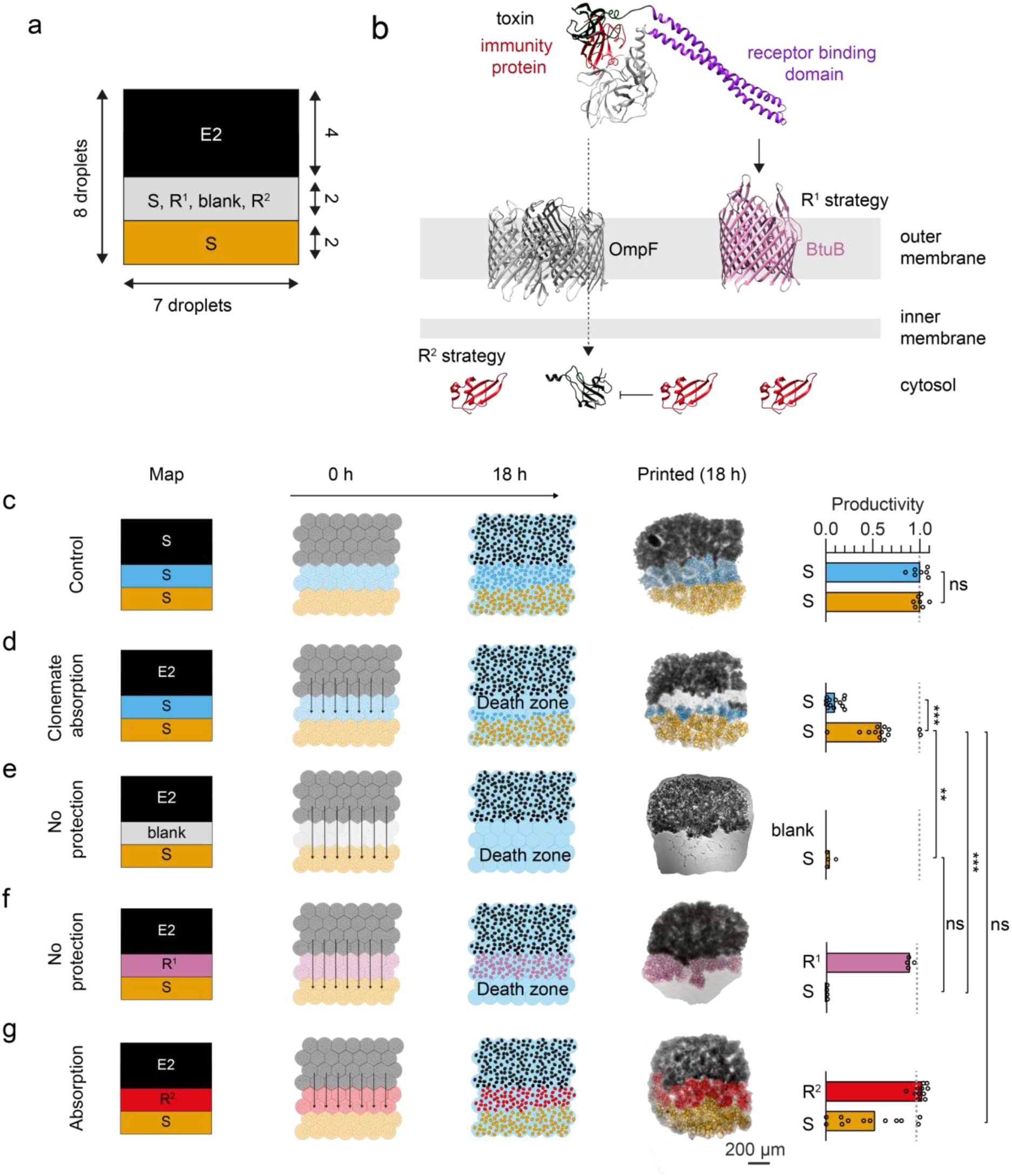
Susceptible strains can provide protection to clonemates against producer strains. **a**, Schematic of the printing map used for confirming clonemate absorption. The producer strip (E2) consists of 7 x 4 x 1 droplets (black), the changeable strip consists of 7 x 2 x 1 droplets (grey), and the sensitive strip consists of 7 x 2 x 1 droplets (orange) in the *x, y*, and *z* dimensions, respectively. **b**, A schematic of the two resistance strategies used for creating R^1^ and R^2^. R^1^ is resistant to colicins because of deletion of the BtuB receptor (which normally binds colicins). R^2^ is resistant to colicins through constitutive expression of the cognate immunity protein, which binds and inactivates the toxin when it enters the cytosol. **c–g**, Printing maps, competition schematics, composite brightfield and fluorescence microscopy images, and bar charts of printed communities comprising three bioinks. **c**, The control printed arrays containing three S strains (*n* = 7). A Wilcoxon test showed the median productivities of S-blue (S *rfp*) and S-orange (S *gfp*) were non-significantly different (P > 0.9999, see Supplementary Table 5 for statistical test). **d**, Printed arrays containing E2 cells (black) and two S strains (*n* = 13). A Wilcoxon test showed the median productivities of S-blue and S-orange were significantly different (P = 0.0002, see Supplementary Table 6 for statistical test). **e**, Printed arrays containing E2 cells, a strip of agarose not containing bacteria, and S cells (*n* = 4). Mann Whitney tests showed the median productivities of S-orange in **e** and **d**, and **e** and **f** were significantly (P = 0.0029) and non-significantly (P = 0.1143) different, respectively (see Supplementary Tables 7 for statistical test). **f**, Printed arrays containing E2 cells, R^1^ cells (reddish purple) and S cells (*n* = 4). A Mann Whitney test showed the median productivities of S-orange in **d** and **f** were significantly different (P = 0.008, see Supplementary Table 8). **g**, Printed arrays containing E2 cells, R^2^ cells (red), and S cells (*n* = 12). A Mann Whitney test showed the median productivities of S-orange in **d** and **g** were non-significantly different (P = 0.604, see Supplementary Table 9). In **d–g**, arrows on schematics represent colicin reach during the competition experiments (not to scale). In the bar charts, each data point is a biological replicate and the height of the bars are mean productivity values. In **c–g**, the term ‘productivity’ (*p*) of a genotype *a* in a printed array after 18 h of competition is defined as: *p* = *A_a_*/*A_s_*, where *A_s_* is the total cross-sectional area that a susceptible genotype occupies after 18 h of competition (at the same starting density and spatial pattern as genotype *a*), but without interference competition.

The ecological interactions within microbial communities are central to their properties and impacts^1,7^. However, the fine spatial scales at which microbes position and interact is a challenge for experimental microbiology. Here we have presented a new method for arranging microbial genotypes in defined 2D or 3D arrays with micron-scale resolution. This method allowed us to establish a causal link between the arrangement of bacterial strains at these fine scales and ecological outcomes. We find that spatial structure impacts the outcome of both interference and exploitative competition. Moreover, fine-scale structure can shift the outcome of competition from one strain dominating to another, and it can be difference between one strain thriving and mutual destruction. A key principle that emerges across our experiments is that spatial segregation is often protective and limits the effects of one strain on the other. We show that for colicin E2, this is explained by the ability of susceptible cells to bind the toxin rather than by diffusion limitation. This finding fits well with a large body of work on bacterial biofilms, which emphasises how both the binding and breakdown of solutes can generate strong gradients over small spatial scales, even when diffusion is not itself limiting^33,34,53–56^. The commonness of these micron-scale solute gradients suggests that the effects we have documented here will be important across many species and ecological interactions. The manipulation of fine-scale spatial structure, therefore, has significant potential for both understanding and controlling microbial communities.

## Methods

### Preparing aqueous phases

Phosphate-buffered saline (PBS) (1X, pH 7.4, Gibco) was used as purchased. Lysogeny broth Miller (LB-Miller) medium was prepared by adding 10 g tryptone (BD Difco^™^), 5 g of yeast extract (BD Difco^™^) and 10 g of sodium chloride (Fischer) to 1 L of Milli-Q^®^ water and autoclaving. 5X M9 medium (Sigma-Aldrich) without a carbon source (12.8 g L^-1^ Na_2_HPO_4_.7H_2_O, 3 g L^-1^ KH_2_PO_4_, 0.5 g L^-1^ NaCl, 1 g L^-1^ NH_4_Cl, 2 mM MgSO_4_, 0.1 mM CaCl_2_) was dissolved in Milli-Q^®^ water, autoclaved, and filtered through 0.22 μm polyethersulfone membrane (Millex-GP Syringe Filter Unit). LB-Miller agar solutions (molten) were prepared by combining LB-Miller medium with agar (final concentration of 1.5% w/v) (BD Difco^™^) and autoclaving. 2.0% w/v of ultra-low-gelling-temperature (ULGT) agarose (Sigma-Aldrich) was prepared by dissolving 100 mg in 5 mL of LB-Miller medium and autoclaving. The hydrogel was kept molten by keeping the solution in a static incubator at 37°C. Molten solutions were only used for a week. Antibiotics were dissolved in Milli-Q^®^ water, filter-sterilised (0.22 μm polyethersulfone membrane), and frozen (−20°C) as stock solutions at 100 mg mL^-1^ for ampicillin and 30 mg mL^-1^ for kanamycin (Sigma-Aldrich).

### Preparing lipid/oil solutions

Lipids were purchased from Avanti^®^ Polar Lipids in powder form and stored at −80°C. Undecane and silicone oil AR20 (Sigma-Aldrich) were filtered before use through 0.22 μm filters (Corning^®^) under vacuum. An optimised lipid/oil solution was prepared for printing^57^. Lipid films were prepared by bringing ampoules to room temperature and dissolving in anhydrous chloroform (Sigma-Aldrich) to make a chloroform/lipid solution at lipid concentrations of 25 mg mL^-1^. Chloroform/lipid stock solutions were used immediately. Using glass syringes (Hamilton^®^), 45 μL and 20 μL of lipid stock solutions 1,2-diphytanoyl-sn-glycero-3-phosphocholine (DPhPC) and 1-palmitoyl-2-oleoyl-sn-glycero-3-phosphocholine (POPC) stock solutions, respectively, were transferred into a 7-mL, isopropanol-cleaned, Teflon-capped glass vial (Supleco^®^). The lipid/chloroform solution was evaporated under a slow stream of nitrogen while rotating by hand to produce an even lipid film. The film was dried under vacuum for 24 h, then stored under house nitrogen at −80°C until use. When required for printing, films were left at room temperate for 30 min, then 2 mL of a pre-mixed solution of undecane and silicone oil AR20 (35:65 by volume) was added to the film, followed by sonication (Branson 2800 ultrasonic bath 230 V) for 1 h. The total concentration of lipids was 1 mM with a molar ratio of DPhPC:POPC of 2:1. Lipid films were kept for a maximum of two months.

### Construction of recombinant DNA

Plasmid and primers used in this study are listed in Extended Table 1 and 3, respectively. All PCRs were performed using KOD Hot Start DNA polymerase (Novagen^®^), following standard protocols. Plasmids were assembled using one-step isothermal assembly (NEBuilder^®^ HiFi DNA Assembly Master Mix M5520AA). All constructs were sequenced to verify sequences were correct (Source BioScience). The generation of pGRG25-P*max:sfgfp, pGRG25-Pmax:mrfp1*, and pGRG25-Pmax:*immE2* plasmids is described in previous work^42^. The plasmids coding for the fluorescent proteins YPet (pBC11-*PybaJ:ypet*) and mNeonGreen (*pBC43-PybaJ:blFP-Y3*) were provided by Olivier Cunrath. The plasmid coding for the IPTG-inducible expression of ImmE2 (*pC001-Ptrc:immE2*^58^) was provided by Nick Housden and Colin Kleanthous. The generation of the plasmid from which ImmE2 is constitutively expressed (pTML9-P*max*:*immE2*) was assembled from the *pBC22-Pybaj:bfp* backbone (a plasmid identical in sequence to the plasmids encoding the YPet and mNeonGreen fluorescent proteins but expressing BFP) using the primers TML-P57 and TML-P58, the ImmE2 coding sequence from pC001-P*trc*:*immE2* using the primers TML-P61 and TML-P62, and the promoter and RBS driving the expression of superfolder GFP in pGRG25-P*max:sfgfp* using TML-P59 and TML-P60. All the primers used in this study were generated using SnapGene^®^.

### Construction of bacterial strains

Bacterial strains used in this study are outlined in Table 1 and Extended Table 2. *E. coli* BZB1011 and BZB1011 pCol genotypes are from previous work^42,59^. For the construction of chromosomally-labelled (*Tn7* insertion) sfGFP and mRFP1 BZB1011 strains and BZB1011 pCol genotypes, see previous work^42^. The BZB1011 *ΔbtuB sfgfp::Tn7* or *mrfp1::Tn7* strains were created by using the method described by Cianfanelli *et al*^60^. Briefly, the regions (700 base pairs) upstream and downstream of the *btuB* gene were amplified by PCR and joined together in the pFOK plasmid using the NEB^®^ HiFi DNA Assembly Master Mix. 5 μL of the reaction was then transformed into chemically competent *E. coli* Jke201 (recovery in 1 mL LB-Miller medium + 100 μM diamino pumilic acid (DAP) and selected on LB-Miller agar plates (1.5% w/v) supplemented with 100 μM DAP and 50 μg mL^-1^ kanamycin). Transformants were screened by colony PCR (GoTaq G2 polymerase) and sequencing (Source Bioscience) using the primers oOPC-614 and oOPC-615. Three clones carrying the correct plasmid (pTML8) were then conjugated with BZB1011 *sfgfp::Tn7* and BZB1011 *mrfp1::Tn7*. Trans-conjugants for which a second recombination event had occurred were then counter-selected. Clones were finally verified by colony PCR and sequencing (by using the primers TML-P9 and TML-P10) for deletion of the *btuB* gene. The BZB1011 *immE2::Tn7* pColE2 *pBC11-PybaJ:ypet* or *pBC43-PybaJ:blFP-Y3* genotypes were generated by transforming competent BZB1011 *immE2::Tn7* pColE2 cells (created in previous work^42^) with respective plasmids, and by selecting for transformants on LB-Miller agar plates (1.5% w/v) supplemented with 30 μg mL^-1^ of kanamycin. The BZB1011 *mrfp1::Tn7 pC001-Ptrc:immE2* and BZB1011 *mrfp1::Tn7 pTML-9-Pmax:immE2* genotypes were generated by transforming competent BZB1011 *mrfp1::Tn7* with either *pC001-Ptrc:immE2* or pTML-9-P*max*:immE2, and selecting for transformants on LB-Miller agar plates (1.5% w/v) supplemented with 100 μg mL^-1^ of ampicillin

### Competent Cells Preparation and Transformation

The strain for transformation was inoculated in LB-Miller medium overnight. The next day, the optical density at 600 nm (OD_600_) of the overnight culture was measured using a spectrophotometer (Thermo Scientific, Genesys 10S UV-VIS). The overnight culture was then inoculated into 50 mL of LB-Miller medium to a final OD_600_ of 0.05, and then incubated at 37°C (shaking at 225 rpm) until the OD_600_ reached ≈ 0.4. Next, the cultures were split into centrifuge tubes (25 mL in each) and centrifuged at 4°C for 10 min at 1734 g. Following this, the supernatants were discarded, and each pellet was gently resuspended in 12.5 mL of 4°C 100 mM CaCl_2_ + 15% v/v glycerol solutions. After addition of these solutions, cultures were left on ice for 45 min. After this, cultures were again centrifuged at 4°C for 10 min at 1734 g. Centrifuged pellets were recombined pairwise by resuspending one of the pellets in 5 mL of ice-cold 100 mM CaCl2 + 15% v/v glycerol solution, and using this resuspension, the second pellet was resuspended. Following this, 200 μL of competent bacteria were aliquoted into 1.5 mL centrifuge tubes and directly frozen at −80°C.

To transform competent cells, 100 μL of competent cells were thawed on ice. Once thawed, DNA (≈100 ng for plasmids or 5 μL of Gibson assembly reaction mixtures) was added to the thawed cells. The cells were then kept on ice for 30 min. Following this, the cells were heat-shocked by transferring the tube containing the cells and DNA into a 42°C water bath for 45 s and then transferring the tube back on ice for 2 min. The cells were then recovered by adding 900 μL LB-Miller medium to the cells and incubating the cells for an hour at 37°C (shaking at 225 rpm). After one hour of recovery, 100 μL of the cultures were plated on selective LB-Miller agar (1.5% w/v). The remaining 900 μL were centrifuged for 1 min at 8000 g, followed by discarding of the supernatant and resuspension of the pellet in 100 μL of LB-Miller medium. The final 100 μL were then plated on corresponding selective LB-Miller agar (1.5% w/v). Both plates were then incubated overnight at 37°C in a static incubator and checked for transformants the following day.

### Growth curves and analysis

Overnight cultures of the strains tested were normalised to an OD_700_ (Optical density taken at 700 nm)^61^ in PBS and washed twice. 10 μL of normalised cultures were added to the wells (containing 190 μL LB-Miller medium) of a flat-bottom transparent 96 well-plate (excluding the two first and two last columns, which were filled with 200 μL of sterile Milli-Q^®^ water), bringing the initial OD_700_ to 0.01. The layout of the plate was randomised using the CRUK PlateLayout tool (https://github.com/crukci-bioinformatics/PlateLayout) for each biological replicate (*n* = 3). Three wells were left uninoculated to calculate background OD700. The plate was then incubated at 37°C (shaking at 400 rpm, double orbital) in a plate reader (BMG Labtech Omega) for 24 h, measuring OD700 every 5 min.

Doubling times were calculated from the growth curves using a custom Python script. In brief, data was imported in Python dataframes, sorted, and the mean of the technical replicates calculated. The mean of the biological replicates was then calculated. The data was then plotted, and the doubling times calculated using the data points between OD 0.2 and 0.7 (exponential phase). Lag phase duration was calculated using DMFit 3.5 (https://www.combase.cc/index.php/en/8-category-en-gb/21-tools)^62^. Plate reader data were formatted to be used by DMFit using a custom Python script.

### Growth Inhibition Assay

Resistance of BZB1011 *sfgfp::Tn7 ΔbtuB*, BZB1011 *mrfp1::Tn7 ΔbtuB*, BZB1011 *mrfp1::Tn7* pC001-P*trc*:*immE2*, and BZB1011 *mrfp1::Tn7* pTML-9-P*max*:*immE2* strains to colicin E2 were assessed using a growth inhibition assay adapted from White *et al*^63^. Each strain to be tested and BZB1011 pColE2 were inoculated in 3 mL of LB-Miller medium and incubated overnight at 37°C (shaking at 225 rpm). The following day, the overnight cultures of the strains to be tested were diluted in LB-Miller medium to an OD600 of 0.05 and incubated at 37°C (shaking at 225 rpm) until they reached an OD600 between 0.6 and 0.7. 200 μL of the cultures were then added to molten soft LB-Miller agar (0.75% w/v) kept at 56°C and poured over LB-Miller agar plates (1.5% w/v). After the plates had dried, 2 μL of serially diluted E2 overnight culture was spotted on the plates. Plates were then incubated at 37°C (static) overnight and imaged to check for inhibition zones or lack of inhibition zones (which determined resistance).

### Preparation of bioinks

Genotypes were cultured directly from glycerol stocks (50% v/v glycerol) in 4 mL of LB-Miller medium for 15 h (shaking at 250 rpm). From the overnight cultures, 20 μL of each was added to 2 mL of fresh LB-Miller medium and cultured for 3 h (shaking at 250 rpm). Cultures were then normalised to 10^9^ cells mL^-1^ by centrifugation at 8000 g for 5 min and resuspended in 100 μL of fresh LB-Miller medium. Centrifugation, removal of the supernatant, and addition of fresh LB-medium (100 μL), were repeated twice to remove any colicin in the supernatant. If 1:1 competitions were investigated at an *SI* of 0.50, the two normalised bacteria cultures were combined at a 1:1 ratio (50 + 50 μL). At any other *SI* (that required two or three nozzles for printing), a bioink was created per culture. To create the bioinks, 150 μL of molten 2.0% w/v ULGT agarose in LB-Miller medium (37 °C), 30 μL of LB-Miller medium, and 20 μL of normalised culture were combined in PCR tubes, and mixed by gently pipetting back and forward using a 200 μL pipette to homogenise the solution for printing. Pipette tips (Corning^®^ 1–200 μL Filtered IsoTip™) and PCR tubes were maintained at 37°C before homogenising the bioink.

### Printing bacterial droplet networks containing bacteria

The functioning of the 3D droplet printer is outlined in previous work^36^. In brief, a custom-built piezoelectric transducer (±130 V) transmits controlled pressure impulses to a chamber filled with Milli-Q^®^ water. The chamber is connected to a glass nozzle (with a tip diameter of ≈ 150 μm) from which the aqueous printing solution is ejected. To prevent mixing of the printing solution with the water inside the chamber and nozzle, an undecane oil plug of ≈5 μL was used to separate the two solutions. A quartz cuvette (Starna Scientific Ltd) with dimensions of 20 x 10 x 10 mm (length, width, and height) was filled with 1 mL of lipid/oil solution and mounted on the micromanipulator. An IR heater (Beurer Ltd., 150 W) was used to equilibrate the temperature of the nozzles and the quartz cuvette (from a fixed distance) to approximately 37°C over 30 min. After equilibration, the bioink was loaded into the nozzle, and the tip was immersed into the lipid/oil solution contained in the quartz cuvette, which was positioned on a digitally controlled micromanipulator (PatchStar micromanipulator, Scientifica, 20 nm resolution). The micromanipulator movements and the piezo actuation were synchronised using a custom-made software developed in LabVIEW (National Instruments™). Printing was monitored using a side-on stereomicroscope (Nikon^®^ SMZ745T) and videos and pictures were acquired using a digital camera (Thorlabs DCC1645C) mounted on the microscope. For creating patterns at *SI* = 0.67, 0.80, 0.88, and 0.94, a second nozzle was used simultaneously to the first. The second nozzle was normalised to the off-set of the first nozzle so the pattern could be printed line-by-line. For three bioinks, the two smaller strips were printed first using two nozzles followed by replacing the first nozzle with a new nozzle containing the third bioink for printing the larger strip. The third nozzle was aligned with a control droplet to complete the pattern.

### Creating printed arrays containing bacteria from printed droplet networks

Once droplet networks were printed, the structures were left for 15 mins at 37°C to allow the droplets to reach their equilibrium contact angles. Following equilibration, networks were moved to the fridge (4°C) for 1 min 30 s to partially gel the ULGT agarose. Next, networks were moved from the fridge to ambient temperature (22°C) and washed with silicone oil AR20 by initially removing 500 μL of the lipid/oil solution and then adding 600 μL of silicone oil AR20. Then, 600 μL of the diluted lipid/oil solution were removed followed by addition of 600 μL silicone oil AR20. This was repeated two more times. The silicone oil washing step was performed to precipitate the lipids so that the tessellated droplets of partially gelled ULGT-agarose could connect with each other. After washing, the networks were left for 5 min at ambient temperature to allow complete breaking of all bilayers and consequent connection of partially gelled ULGT agarose droplets. After 5 mins, the networks were moved to the fridge for 45 min to create a gelled block of LB-Miller ULGT-agarose containing patterned bacteria (printed arrays). After 45 min, each printed array was moved to individual containers filled with 600 μL of LB-Miller media (Nunc^™^, Lab-Tek^™^ II Chamber Slide™ System (8 wells)) by hydrating the array in 0.5 μL of LB-Miller medium (using a 0.5-μL Hamilton^®^ syringe), followed by picking up the LB-Miller droplet containing the printed array (using a cut pipette tip (20 μL)) and pipetting the printed array into the container. Sterile 20 μL pipette tips were cut using a sterile scalpel (No. 21 Swann-Morton^®^). Printed arrays were encapsulated in LB-Miller medium and subsequently transferred into chambers by using a Leica^®^ EZ4 Stereomicroscope with a 16X eyepiece.

### Bacterial competition experiments within printed arrays and imaging of results

For competition experiments, printed arrays in LB-Miller media were incubated at 37°C for 18 h. After 18 h, printed arrays were washed with M9 medium to remove excess bacteria that grew out into solution from the printed arrays during the competition experiment. Within each compartment containing a printed array, 400 μL of LB-Miller medium were removed and 200 μL of M9 medium were added. Following this, 200 μL of the medium were removed and another 200 μL of M9 medium were added to further wash and remove the bacteria. Finally, printed arrays were picked up using a cut pipette tip (20 μL) and transferred into individual containers containing 400 μL of M9 media (Nunc^™^, Lab-Tek^™^ II Chambered Coverglass (8 well)). Printed arrays were left in M9 media for 10 hours to allow exchange of LB medium with M9 medium, required for imaging. Printed arrays were imaged using a Leica^®^ SP5 confocal microscope using a HC PL Fluotar 10X/0.30 objective (0.3 numerical aperture), at excitation wavelengths of 405 nm and 546 nm and emission cut-offs at 420–520 nm and 625 nm, respectively (exciting superfolder GFP, mRFP1, YPet, and mNeonGreen), a *z*-step of 2.73 μm, and scanning speed of 400 Hz.

### Colony-forming units of printed arrays

Printed arrays were washed and transferred to 400 μL of M9 medium after 8 h of competition. This timepoint was chosen because space in the printed arrays was saturated with *E. coli* within 8 h. Next, printed arrays were moved to individual 1 mL centrifuge tubes containing 200 μL of M9 medium and melted at 55°C for 1 min in a thermomixer (Eppendorf ThermoMixer^®^ C) shaking at 1000 rpm. Following this melting, the *E. coli*-containing solutions were serially diluted by 10 000-fold (100 μL in 900 μL of M9 medium). Following this, 10 000X solutions were plated by beads on 1.5% w/v LB-Miller agar plates, and then left to grow for 24 hours at 37°C in a static incubator. After 24 hours, colonies were counted using a Zeiss PlanApo Z 0.5X objective on an AxioZoom.V16 Zeiss microscope. Genotypes were recognised by fluorescence.

### Flow cytometry of printed arrays

Printed arrays were washed and transferred to 400 μL of M9 medium after 18 h of competition. Next, printed arrays were moved to individual 1 mL centrifuge tubes containing 200 μL of M9 medium and melted. The cell suspensions were diluted a further 10-fold in M9 medium. Diluted cell suspensions were analysed on a BD Accuri C6 Flow Cytometer using 10 000 and 8 000 as thresholds for the forward and side scatter parameters, respectively. For each experimental condition, 20 000 events were quantified per sample. Data was analysed using a custom written script in R. The population of bacteria was gated by considering only the events within the core of the cell population in order to minimize count biases (Supplementary Fig. 1c). This was calculated from the forward and side scatter dimensions. Cells that had a fluorescence intensity above 2^9.5^ were assigned as BZB1011 *sfgfp::Tn7*. Cells below this were assigned as BZB1011 *mrfp1::Tn7* (Supplementary Fig. 1d). Counts shown in Extended Fig. 2c are based on the first 3000 cells that met the selection criteria, which was used to calculate the frequencies of BZB1011 *sfgfp::Tn7* and BZB1011 *mrfp1::Tn7*.

### Microcolony quantification by image analysis

2D segmentation: 3D confocal images of printed arrays were projected into 2D using an average intensity projection (using FIJI^64^ software). The resulting 2D images were then segmented using a combination of ridge detection^65^, morphological watershed^66^, and intensity thresholding. This generated binary images of microcolonies. Each binary image was then used as a mask to extract each microcolony’s position, orientation, length and width, as well as its brightness in the two fluorescence channels. Image segmentation and analysis were performed using MATLAB^®^ (MathWorks^®^) with a custom-written generated user interface (*FAST* – http://doi.org/10.5281/zenodo.3630642).

Print isolation: Some images of prints contained small pieces of detached debris and regions from other prints. To automatically remove these from downstream analyses, the size of each print was estimated and used to exclude these external microcolonies. In the first step of this process, the approximate centre of the print 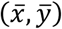 was found by averaging the *x* and *y* coordinates of all microcolonies in the image. The average distance of each microcolony from 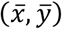 was then calculated as 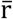. The approximate side length of the print L was then found as:

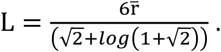

This was used to set a radial distance threshold on the detected microcolony positions, allowing the main print to be isolated from unrelated microcolonies. 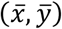 was then recalculated excluding these microcolonies, providing a more accurate measure of the centre of the print. Radial distributions were calculated using this position as their centre point.

Print registration: The above approximation of L assumes that prints are perfectly square. To obtain a more accurate measure of the size of the rectangular prints, they were first registered to the *x-y* coordinate system. This required finding an angle θ through which each print could be rotated to bring it into alignment with all other prints. The first stage of this process involved finding the convex hull of the print, and then measuring the distance between each point on the hull and 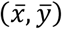. These values were then resampled onto a regularly spaced series of *N* polar angles *φ_n_* measured from 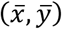. The mean was then subtracted from the resulting series of values, resulting in a function *d*(*φ*) that was positive at values of *φ* where the outer boundary of the print was further-than-average away from the centre and negative at values of *φ* where it was closer than average.

*d*(*φ*) has peaks at values of *φ* corresponding to each of the four corners of the print interleaved with troughs at values of *φ* corresponding to the sides. Furthermore, *d*(*φ*) is circular, with *d*(*φ*_1_) = *d*(*φ*_*N*+1_). θ could therefore be found from the phase of the Discrete Fourier Transform (DFT) on *d*(*φ*) with wavenumber *k* = 4. Explicitly,

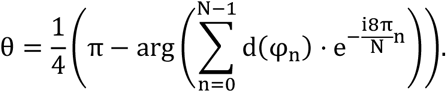

The position of each microcolony was then rotated about 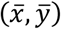 by θ, and the size of the print in each direction estimated as the difference between the minimum and maximum *x* and *y* coordinates of the registered microcolonies.

Microcolony neighbour-neighbour distance estimation: Delaunay triangulation^67^ was applied to the positions of all microcolonies in the print. Microcolonies directly connected in the resulting triangulation were assigned as neighbours. Due to small undulations in the shape of the print, some microcolonies located at the outer boundary of the print were assigned to very distant neighbours. Microcolonies forming the outer boundary of the print were therefore excluded when calculating the radial neighbour-neighbour distance profile (Extended Fig. 2g).

3D segmentation: 3D segmentation of confocal microcolony data was performed using a generalisation of the algorithm used during 2D segmentation. Specifically, a rough segmentation of the initial array of voxel intensities *I*(*x,y,z*) was first generated by applying a simple intensity threshold, allowing the bright microcolony-containing regions to be separated from the dark background. However, variations in background intensity and intermediate intensities of voxels between closely-packed microcolonies prevented this approach from reliably isolating individual microcolonies within these regions.

To separate these closely-packed microcolonies, we located 3D ridge-like features (e.g. blobs, sheets or tubes) that were dark relative to nearby microcolonies^68,69^. Initially, *I*(*x,y,z*) was convolved with a Gaussian kernel *G*(*x, y, z, σ*) of spatial scale *σ* to isolate features at that scale:

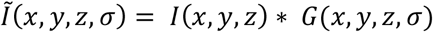

The second-order derivatives of *Ĩ*(*Ĩ_xx_, Ĩ_xy_* etc.) were next used to construct the Hessian *H* of the image at each voxel:

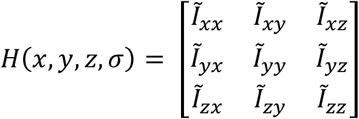

*H* contains information on the 3D features of scale *σ* present within the image around position (*x,y,z*). These features can be classified by finding the values of the eigenvalues *λ*_1_, *λ*_2_ and *λ*_3_ of *H*, where |*λ*_1_| ≥ |*λ*_2_| ≥ |*λ*_3_|^68,69^. However, as we were not concerned with the specific feature separating microcolonies, we generated a binary segmentation of the image simply by finding all voxels for which *λ*_1_ was above a threshold value. This fine-grained mask was then combined with the initial rough segmentation using a voxelwise AND filter, generating the final segmentation.

### Calculating segregation indices

We used a segregation index (*SI*) to measure genetic mixing within printed arrays, and to quantify printing fidelity. *SI* were computed for printed-colony micrographs as follows.

1. Segmentation. Each micrograph pixel was assigned a genetic identity (‘A’, ‘B’ or ‘None’) by comparing its two fluorescence channel intensities, thresholded separately using Otsu’s method^70^. Pixels in which both channels exceeded the intensity threshold were classified as containing neither cell type (‘None’).
2. Local *SI*. We computed a local segregation index *s_i_* for each A- or B-type pixel *i*, by comparing its genetic identity with those of neighbouring pixels. For a square neighbourhood of size *h, s_i_*(*h*) is defined as

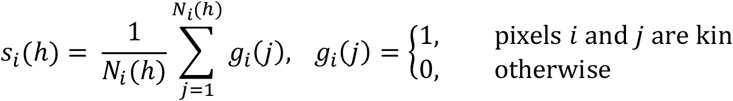

where *N_i_*(*h*) is the number of A- or B-type pixels within *h* rows or columns of focal pixel *i*. Thus, *s_i_*(*h*) is the fraction of *i*’s neighbours that share its genotype. Note: for ease of image processing, here we are considering square neighbourhoods instead of the circular ones used previously^19,71^.
3. Global *SI*. We defined the global segregation index *SI* for a community micrograph as the arithmetic mean of local segregation indices for each of *M* focal pixels,

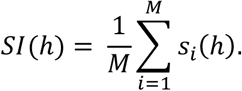 *SI*(*h*) varies in value from 0.5 (high genetic mixing on spatial scale *h*) to 1.0 (no genetic mixing).

To assess printing fidelity, we generated reference images representing idealised community structure (Extended Fig. 2e) and compared the resulting *SI*(*h*) values with those measured for printed communities (Fig. 1h). This comparison revealed a good fit between measured and reference *SI*(*h*) values for a range of spatial scales (Fig. 1h; *h* = 5, 10, 20, 50, 100 px, equivalent to 7.6, 15.2, 30.3, 75.8, 151.5 μm), confirming that community printing is capable of modulating genetic mixing. Image processing and *SI* calculations were carried out using MATLAB^®^ (MathWork^®^).

### Definitions and calculations

The term ‘frequency’ (*f*) of a genotype, *a*, in a printed array containing genotype *a* and *b* after 18 h of competition is defined as:

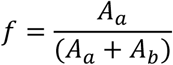

*A* is the total cross-sectional area that a genotype occupies in the printed array. *A* is calculated by segmentation of *z*-projected (by average intensity) confocal microscopy images of the printed array. *f* is used in Fig. 1, 2, and 3. In Extended Figures 2a, *A* = total volume occupied by a strain, number of cells, and number of CFUs; for 3D segmentation, flow cytometry, and CFU, respectively.

The term ‘productivity’ (*p*) of a genotype, *a*, in a printed array after 18 hours of competition is defined as:

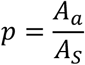

*A_s_* is the total cross-sectional area that a susceptible genotype occupies after 18 h of competition (at the same starting density and spatial pattern as genotype *a*), but without interference competition.

For competition between two genotypes (Fig. 3), this would be the total cross-sectional area occupied by a susceptible strain at the given *SI*, competing against another susceptible strain. For competition between three genotypes (Fig. 4), this would be the total cross-sectional area occupied by a susceptible genotype in the strip (Fig. 4c), competing against the other two susceptible genotypes in the other strips.

## Data availability

All relevant data are available from the corresponding authors upon request.

## Code availability

Our written code used for 2D and 3D segmentation, and sub-sequent analysis of microcolonies of z-projected and z-stacked confocal images of printed arrays is available at: https://zenodo.org/record/3630642#.Xv7ryp5KiEs. Further scripts used to analyse data are available from the corresponding authors upon request.

## Acknowledgements

This research was supported by a European Research Council Advanced Grant SYNTISU, which supports RKK, LZ, GSP, and HB; and by a European Research Council Advanced Grant 787932 and a Wellcome Trust Investigator award 209397/Z/17/Z to KRF. TAM-L and AA acknowledge funding from the University of Oxford, the EPSRC & BBSRC Centre for Doctoral Training in Synthetic Biology (grant EP/L016494/1), and a Clarendon Fund Scholarship. OJM acknowledges funding from the EPSRC Life Sciences Interface Centre for Doctoral Training (grant EP/F500394/1). DG acknowledges funding from the Swiss National Science Foundation Postdoc Mobility Fellowships P2LAP3_155109 and P300PA_167703. DAIM acknowledges funding from the MRC Career Development Award MR/M009505/1. The authors thank the electronic workshop in the Physical and Theoretical Chemistry Department for aid in the design and construction of the piezo drivers; Olivier Cunrath, Nicholas Housden, and Colin Kleanthous for plasmids; and Connor Sharp for useful discussions.

## Author contribution

RKK, DG, WK, GSP, HB, and KRF conceived and designed experiments. RKK and KRF wrote the paper, with comments from other authors. RKK, TAM-L, and AA performed experiments. OJM and WPJS wrote custom software for image analysis of printed arrays and calculation of segregation indices, respectively. DAIM and DG designed and generated the colicin toolbox. LZ designed and built the high-power piezo driver.

## Notes

### Competing Interest Statement

The authors have declared no competing interest.

