## Supplementary Information for "Droplet printing reveals the importance of micron-scale structure for bacterial ecology"

#### Extended Figures

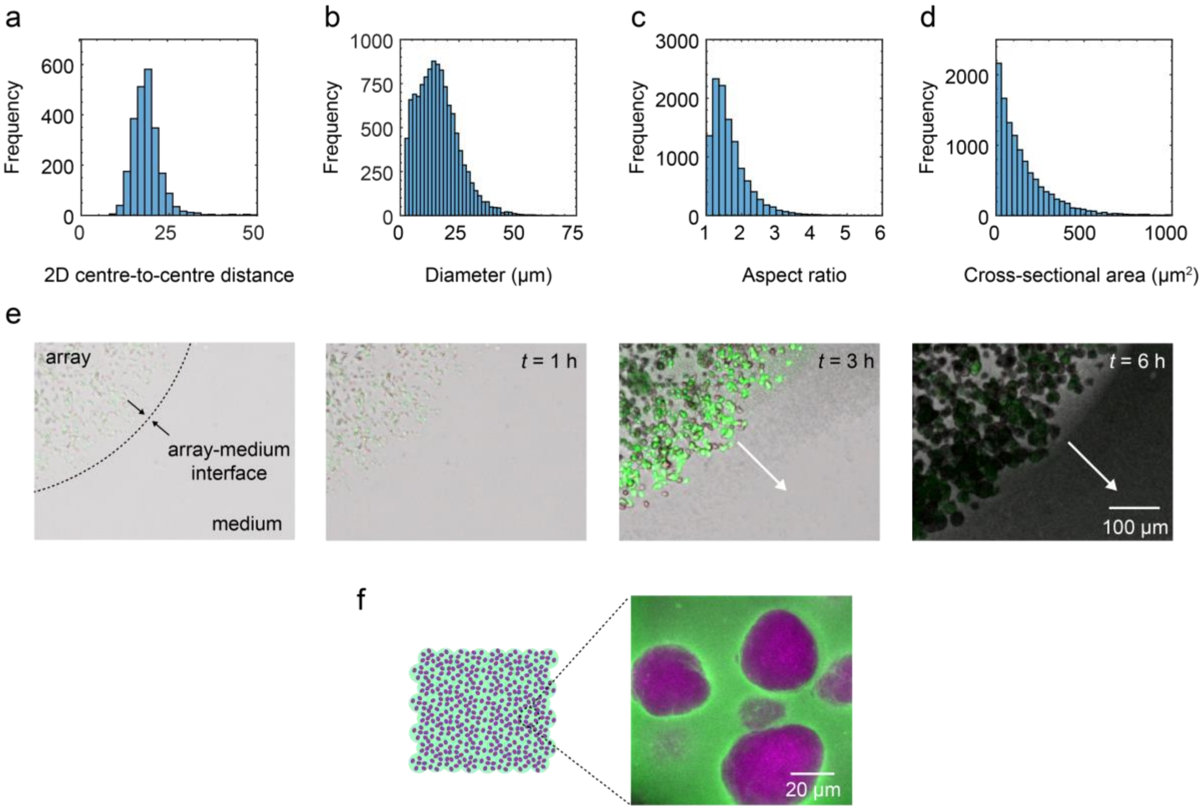


**Extended Fig. 1. Characterisation of the growth of *E. coli* within the printed arrays**. **a**, Histogram of the 2D centre-to-centre distances between *E. coli* microcolonies in the printed arrays. Distances are calculated over $n$ = 6 printed arrays containing susceptible (S) *gfp* and S *rfp* cells printed at an initial *SI* = 0.50 (genetically-mixed) after 18 h of competition. **b–d**, Histograms of microcolony dimensions including the cross-sectional diameter (longest length) (**b**), cross-sectional aspect ratio (**c**), and cross-sectional area (**d**), in $n$ = 6 printed arrays containing S *gfp* and S *rfp* printed at an initial *SI* = 0.50 after 18 h of competition. In **a**–**d**, values are measured from *z*-projected (by average intensity) confocal stacks of printed arrays. **e**, Time-lapse confocal microscopy of a printed array containing S *gfp* and S *rfp* cells printed at an initial *SI* = 0.50. Confocal images are composite *z*-projection images from 20 μm slices. Arrows indicate invasion of *E. coli* from the microcolonies within the printed arrays into the surrounding medium. **f**, A single-slice confocal microscopy image of microcolonies in printed arrays. Magenta microcolonies are false-coloured S *rfp* colonies and green represents fluorescent agarose. Images are taken from printed arrays containing only S *rfp* cells after 18 hours of growth. See Supplementary Methods for further discussion.

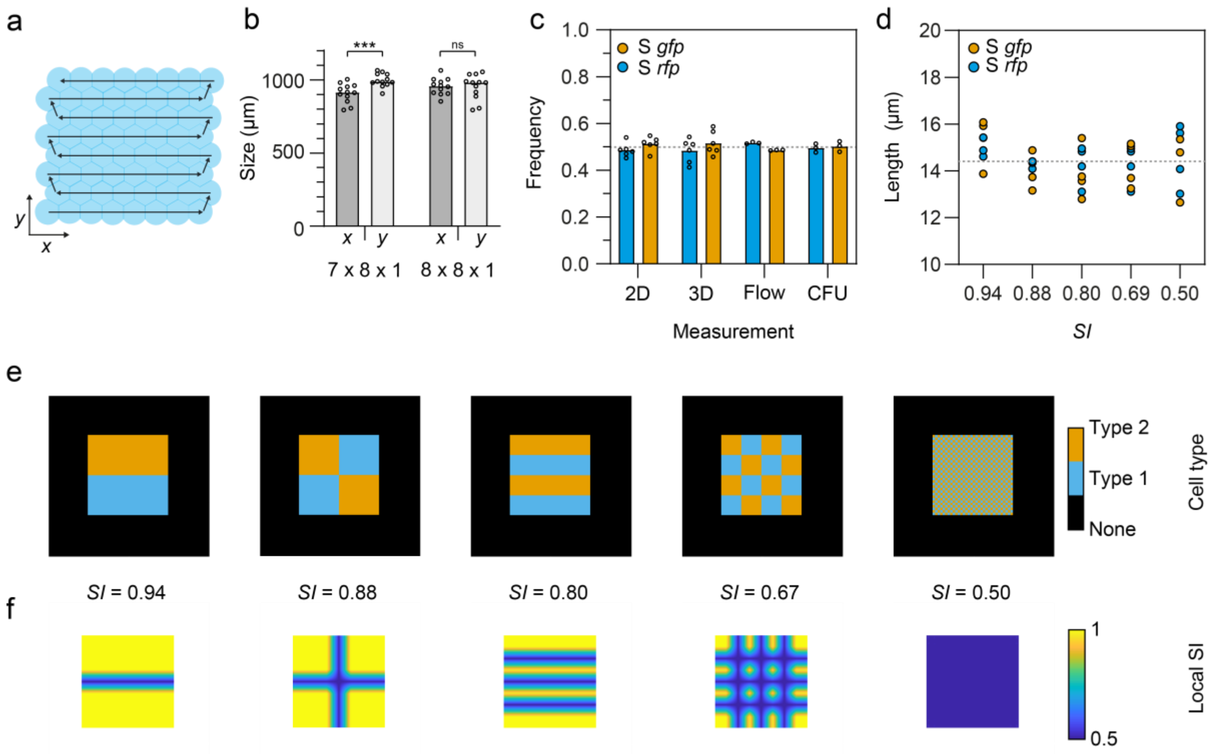


**Extended Fig. 2. Measuring printing fidelity**. **a**, A diagram showing the *x* and *y* dimensions, and the printing path used to generate printed array. **b**, A bar chart showing the 2D sizes (with axes shown in **a**) of printed arrays generated from droplet networks comprising 7 x 8 x 1 droplets (*x*, *y*, *z*), and 8 x 8 x 1 droplets (*x*, *y*, *z*). Each data point represents a biological replicate and the height of the bars are the mean values. For 7 x 8 x 1 and 8 x 8 x 1 printed arrays, unpaired t tests with Welch's corrections showed a significant and non-significant difference between the means of the *x* and *y* dimensions, respectively (see Supplementary Tables 10 for statistical tests). These experiments were performed to assess the size of the printed arrays after printing. The results show that the arrays retain anisotropy if one axis is larger than the other (7 x 8 x 1 arrays), that they swell in medium (as they are formed from 110 µm (in diameter) droplets), and the arrays are sub-millimetre in size. **c**, A bar chart showing the frequency (proportion of cells in the population) of susceptible (S) *gfp* and S *rfp* cells as measured from the total cross-sectional area each strain occupies in a printed array (calculated by 2D segmentation of *z*-projections of confocal stacks (2D)) (*n* = 6), the total volume each strain occupies in a printed array (calculated by 3D segmentation of confocal stacks (3D)) (*n* = 6), flow cytometry (flow) (*n* = 3), and colony-forming units (CFU) by plating (*n* = 3), in arrays printed at an initial *SI* = 0.50 after 18 h of competition. Each data point represents a biological replicate and the heights of the bars are the mean frequencies. A Kruskal-Wallis test found a non-significant difference between the median frequencies of S *gfp* (P = 0.2415; see Supplementary Table 11 for statistical test). This experiment was performed to assess if there was a difference between measurements used to calculate strain-frequencies in printed arrays. As there is a non-significant difference between measurements, we used 2D segmentation to calculate the frequencies of strains in our competition experiments (but see Extended figure 3i further comparisons). **d**, A Graph assessing the printing fidelity of arrays containing S *gfp* (orange) and S *rfp* (blue) cells after 18 h of competition. Plots show the 2D morphologies (lengths) of S *gfp* and S *rfp* microcolonies (from *z*-projected confocal stacks) within arrays printed at different *SI* values. Each data point is a mean value from a different printed array. The grey line represents the mean of the mean lengths. A Kruskal-Wallis test found a non-significant difference between the median lengths of S *gfp* microcolonies at different *SI* values (P = 0.4247; see Supplementary Table 12 for statistical test). These experiments demonstrate that the morphologies of microcolonies were not affected by imposed printing patterns (initial *SI* values)._**e**, Reference images representing idealised community structures. **f**, Representative local *SI* maps of the reference images using a 'span' of 100 µm (see Methods). The structures from left to right have an *SI* value of 0.94. 0.88, 0.80, 0.67, and 0.50, respectively. See Methods for a full description.

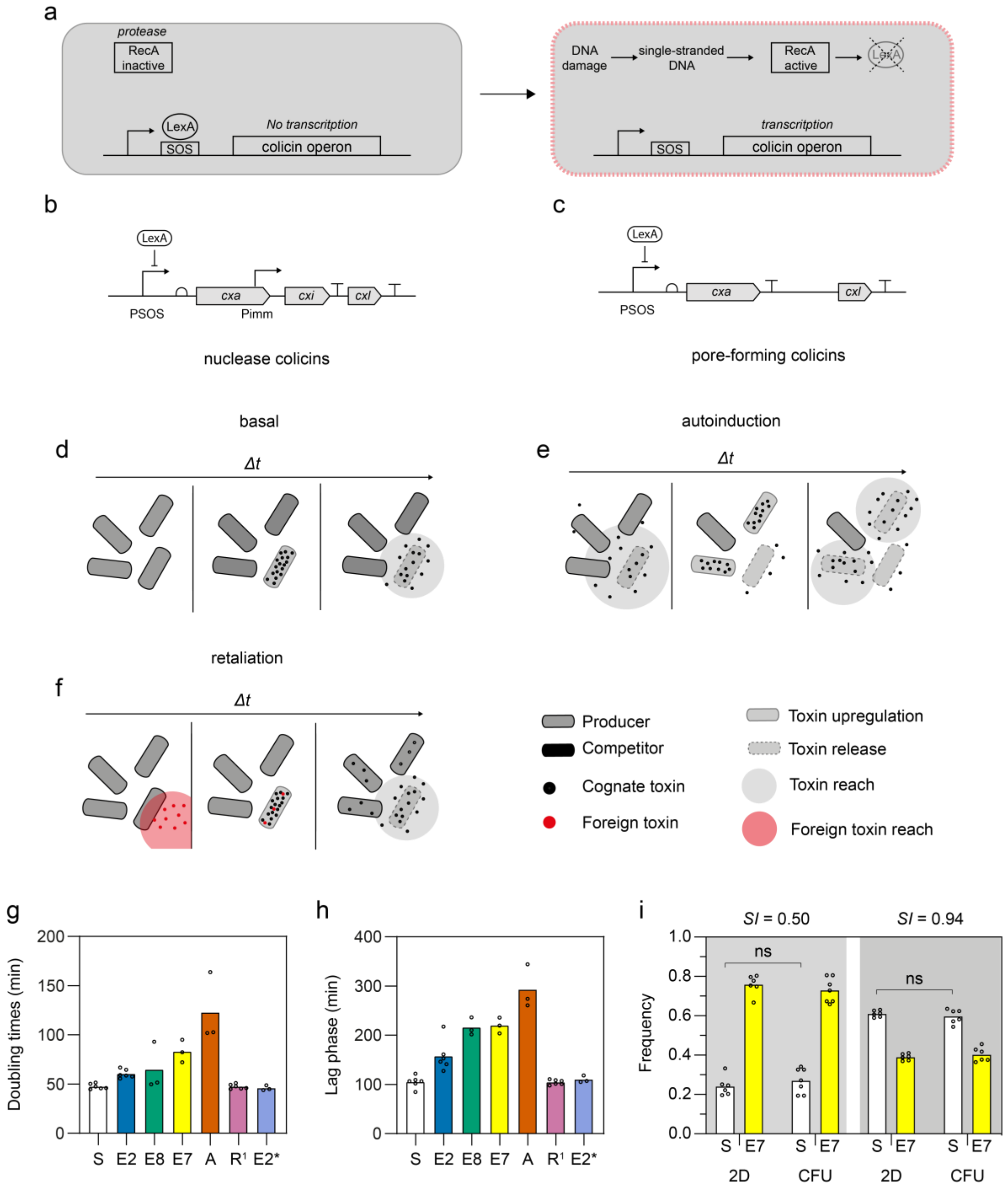


**Extended Fig. 3. Colicin production and growth rates.** **a**, Regulation of colicin production via DNA damage and the SOS response. LexA is the transcriptional repressor and RecA is the protein (upon activation by single-stranded DNA) that activates LexA cleavage. As a result, cells containing the colicin plasmids (pCol) will produce colicins in response to DNA damage. **b**–**c**, Organisation of the colicin operons for nuclease colicins, E2, E7, and E8 (**b**), and the pore-forming colicin A (**c**). The genes *cxa, cxi* and *cxl* are the colicin, immunity, and lysis genes, respectively. *x* denotes the specific colicin and diagrams are modified from [52]. **d**, Basal: colicin production occurs at a baseline level, such that producers will kill susceptible strains without SOS response activation (Fig. 2). **e**, Autoinduction: some strains display autoinduction of colicin production (E2 and E8 here). In these strains, the import of a colicin from a clonemate may activate the SOS response, which means that high cell density of the strain in question amplifies its colicin production (Fig. 2 and 3). **f**, Retaliation: a colicin-producer retaliates when a non-cognate (foreign) toxin damages their DNA. This DNA damage induces the SOS response and consequent expression of the colicin operon. Colicin-producers E7, E2, and E8, antagonise one another because they all produce DNA-damaging toxins. Colicin-producer A retaliates to E2, E7, and E8, but does not cause these strains to retaliate because colicin A targets the membrane, not DNA. This leads to an advantage for A seen in competition with E2 (Fig. 3b). **g**, **h** Bar graphs of doubling times and lag phases (measured from shaken liquid cultures) of strains used in competition experiments. Data points are biological replicates and the heights of the bars are the mean doubling times and lag phase durations. S includes BZB1011 *sfgfp*::*Tn7* and BZB1011 *mrfp1*::*Tn7*, E2 includes BZB1011 *sfgfp*::*Tn7* pColE2 and BZB1011 *mrfp1*::*Tn7* pColE2, E8 is BZB1011 *mrfp1*::*Tn7*, E7 is BZB1011 *mrfp1*::*Tn7*, A is BZB1011 *mrfp1*::*Tn7* pColA, $R^1$ includes BZB1011 *sfgfp*::*Tn7 ΔbtuB* and BZB1011 *mrfp1*::*Tn7*, and E2* is BZB1011 *mrfp1*::*Tn7* pTML9-P*max:immE2*. **i**, Frequencies of S (white) and E7 (yellow) in arrays printed at an initial *SI* = 0.50 or *SI* = 0.94 after 18 h of competition; where frequencies are calculated from the total cross-sectional area a strain occupies in a printed array (which is calculated by segmentation of *z*-projected confocal microscopy images (2D)) or by colony-forming units (CFU) (see Methods). Each data point is a biological replicate and the heights of the bars are the mean frequencies. Mann-Whitney tests showed non-significant differences between median frequencies of S calculated from segmentation and CFU (P = 0.7308 and P = 0.5887 for *SI* = 0.50 and *SI* = 0.94, respectively; see Supplementary Tables 13 for statistical tests).

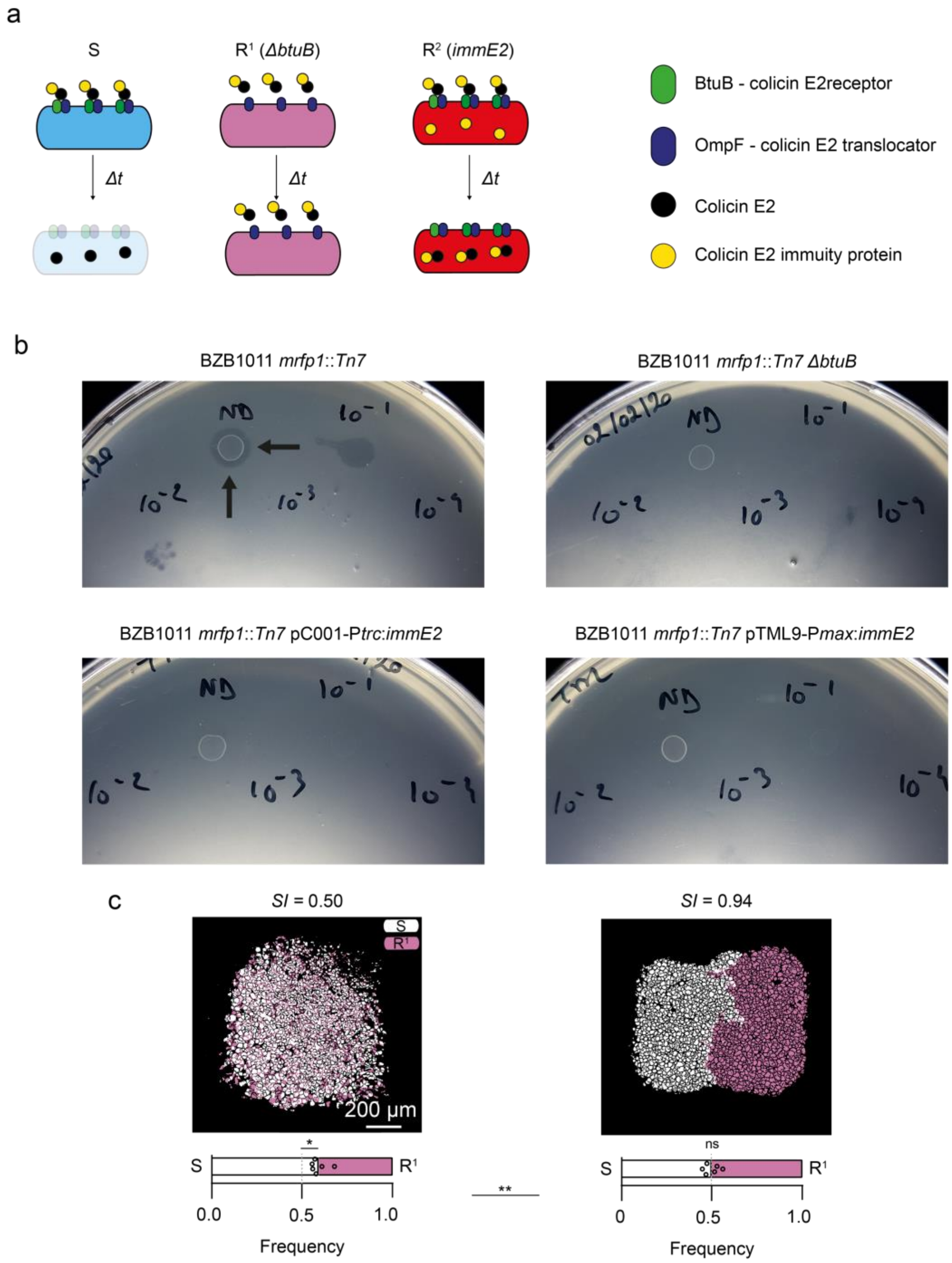


**Extended Fig. 4. Competition with colicin-resistant strains**. **a**, Interaction of colicin E2 with susceptible cells (S), and resistant cells made by *btuB* deletion (R[1]), and resistance cells that overexpress the colicin immunity protein (*immE2*, R[2]). **b**, Images of LB-Miller agar plates for assessing resistance to colicin E2 for BZB1011 *mrfp1*::*Tn7* (S), BZB1011 *mrfp1*::*Tn7 ΔbtuB* (R[1]), BZB1011 *mrfp1*::*Tn7* pC001-P*trc:immE2* (R[2]) without IPTG induction, and BZB1011 *mrfp1*::*Tn7* pTML9-P*max*:*immE2* (R[2]). Inhibition is only seen for BZB1011 *mrfp1*::*Tn7* as indicated by the two arrows. In **b**, BZB1011 pColE2 was spotted on top of the strains of interest without dilution (ND), and after dilution, by $10^1$, $10^2$, $10^3$, and $10^4$-fold. See Methods for growth inhibition assays. These experiments were performed to show that the genetic modification did indeed produce the phenotypic resistance to colicin E2. **c**, Competition between S (white) and $R^1$ (pink) in arrays printed at an initial *SI* = 0.50 and 0.94 after 18 h of competition, which includes segmented fluorescence images and corresponding bar charts of strain frequencies in the arrays. In the bar charts, data points are biological replicates and where the bars meet are the mean strain frequencies. One sample Wilcoxon tests showed a statistically significant and non-significant difference between the median frequencies of S and $R^1$ in arrays printed at *SI* = 0.50 (P = 0.0312) and *SI* =0.94 (P = 0.8438), respectively (see Supplementary Tables 14 for statistical tests). An unpaired Mann-Whitney test showed a statistically significant difference between the median strain frequencies in S vs $R^1$ arrays printed at *SI* = 0.50 and 0.94 (P = 0.0043, see Supplementary Tables 14 for statistical tests). The experiments in **c** were performed to see the effects of exploitative competition in genetically-mixed (*SI* = 0.50) and fully segregated (*SI* = 0.94) printed arrays.

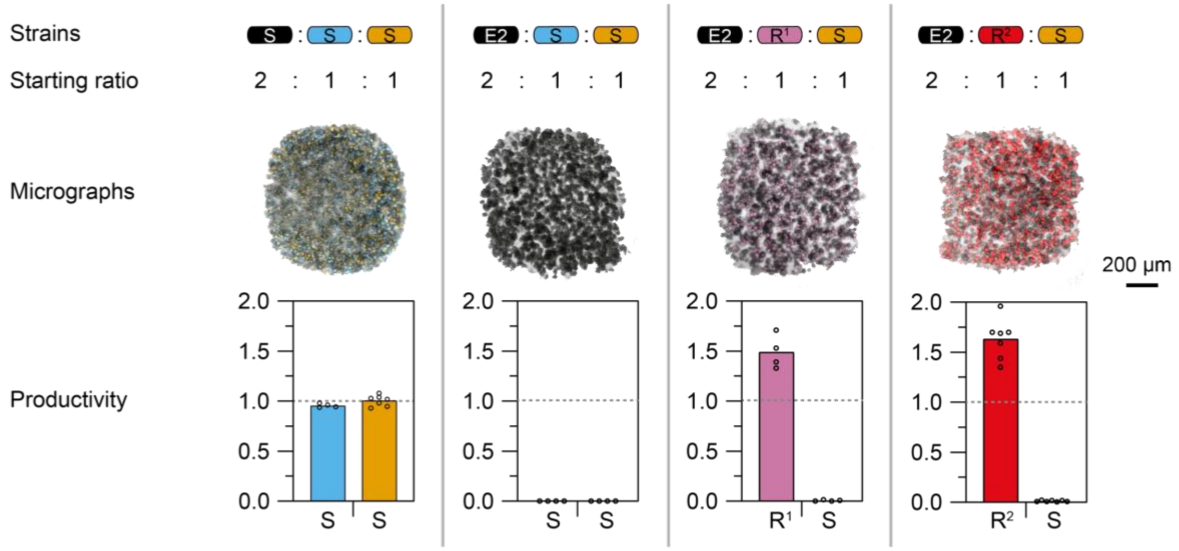


**Extended Fig. 5. The importance of spatial structure for clonemate protection**. Control competition experiments with strain ratios analogous to Fig. 4, but without spatial structure, i.e. all strains are present in each droplet – microscopy images and productivity values are taken and calculated after 18 h of competition, respectively. The goal of this experiment was to confirm that the susceptible strain and its derivatives (R1, R2) are affected as expected by the colicin E2 strain in the absence of spatial structure. Note that we did not follow the productivity of the E2 colicin producers in this experiment, and only show data on the productivity of S, R1, and R2. The microscopy images are composite (including transmitted light and fluorescence) confocal microscopy images. The arrays containing three S strains (black – BZB1011, blue – BZB1011 *mrfp1*::*Tn7*, and orange – BZB1011 *sfgfp::Tn7*) are the genetically-mixed, corresponding experiments to Fig. 4c. The arrays containing E2 (colicin E2 producer) (black – BZB1011 pColE2) and two S strains (blue, and orange) are the genetically-mixed, corresponding experiments to Fig. 4d. The arrays containing E2 (black), $R^1$ (resistant by deletion of the BtuB receptor) (pink), and S cells (orange) are the genetically-mixed, corresponding experiments to Fig. 4f. The arrays containing E2 (black), $R^2$ (resistant by constitutive expression of ImmE2), and S cells (orange) are the genetically-mixed, corresponding experiments to Fig. 4g. In the bar charts, the data points are biological replicates ($n$ = 6 for **a**, $n$ = 4 for **b**, $n$ = 4 for **c**, and $n$ =7 for **d**) and the heights of the bars are the mean productivity values. The grey dashed lines indicate a productivity value of 1.0. These experiments demonstrate that clonemate protection is lost in the absence of spatial structure.

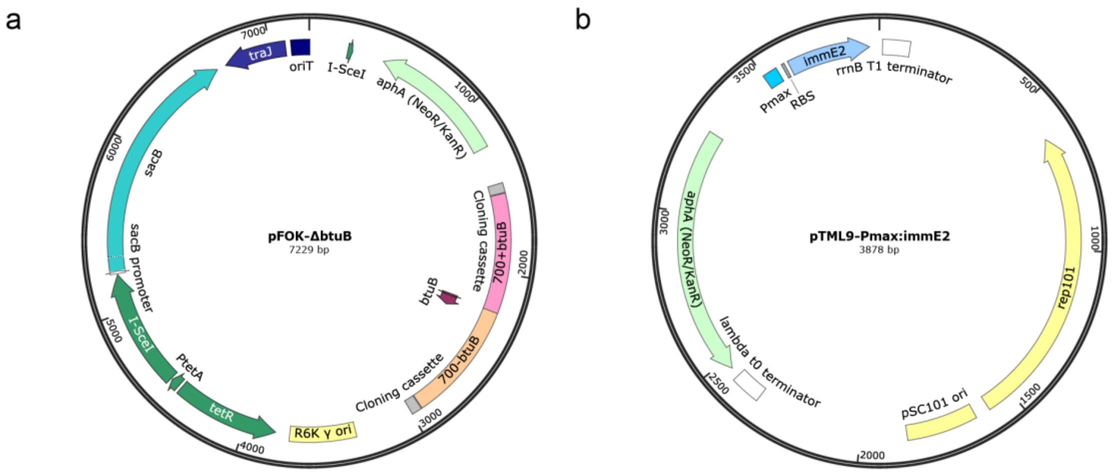


**Extended Fig. 6. a**–**b**, Maps of the pFOK-*ΔbtuB* (**a**) and pTML-P*max*:*immE2* (**b**) plasmids generated in this study.

**Extended Table 1.** Recombinant DNA used in this study.

| Recombinant DNA | Source | Description |
|---|---|---|
| pGRG25-P*max*:*sfgfp* | [1] | Plasmid encoding for superfolder GFP under constitutive expression. Plasmid can integrate at the *Tn7* site on the chromosome by arabinose induction. P*max* is a strong, constitutive BIOFAB promoter. |
| pGRG25-P*max*:*mrfp1* | [1] | Plasmid encoding for mRFP1 under constitutive expression. Plasmid can integrate at the *Tn7* site on the chromosome by arabinose induction. |
| pBC11-P*ybaJ*:*ypet* | Olivier Cunrath | Plasmid encoding for YPet under constitutive expression. P*ybaj* is a constitutive *Salmonella Enterica* promoter. |
| pBC43-P*ybaJ*:*blFP-Y3* | Olivier Cunrath | Plasmid encoding for mNeonGreen under constitutive expression. |
| pBC22-P*ybaj*:*bfp* | Olivier Cunrath | Plasmid encoding for BFP under constitutive expression. |
| pGRG25-P*max*:*immE2* | [1] | Plasmid encoding for ImmE2 under constitutive expression. Plasmid can integrate at the *Tn7* site on the chromosome by arabinose induction. |
| pC001-P*trc*:*immE2* | [2] | Plasmid encoding for ImmE2 under IPTG-inducible expression. |
| pTML9-P*max*:*immE2* | This study (see Extended Fig. 6) | Plasmid encoding for ImmE2 under constitutive expression. |
| pFOK-*ΔbtuB* | This study (see Extended Fig. 6) | Plasmid encoding 700 base pairs of the *btuB* gene. This plasmid was used for deletion of the *btuB* gene. |

**Extended Table 2.** *E. coli* strains and corresponding genotypes used in this study.

| Strain and genotype | Abbreviation | Source | Description |
|---|---|---|---|
| BZB1011 (*λ-*<br>*, gyrA586(NalR), IN(rrnD-rrnE)1, rpsL-(*strR*), rph-1*) | S (Fig. 4) | [3] | Background strain for all genotypes. Susceptible to all colicins. |
| JKE201 (MFDpir *ΔmcrA Δ(mrr-hsdRMS-mcrBC*) *aac(3)IV*::*lacIq)* | na | [4] | Derivative of the MFDpir strain. Lacks EcoKI, the three type IV restriction systems, the restored gentamicin sensitivity, and has the *lacI*$^q$ allele. Used for conjugation of pFOK-*Δbtub* into BZB1011. |
| BZB1011 *sfgfp*::*Tn7* | S *gfp* or S *orange* (Fig. 1,2, and 4) | [1] | Colicin-susceptible strain that constitutively expresses superfolder GFP under P*max* at the *Tn*7 site. |
| BZB1011 m*rfp1*::*Tn7* | S *rfp* or S *blue* (Fig. 1,2, and 4) | [1] | Colicin-susceptible strain that constitutively expresses mRFP1 under P*max* at the *Tn*7 site. |
| BZB1011 *sfgfp*::*Tn7 ΔbtuB* | R[1] | This study | Resistant to group A colicins through deletion of the BtuB receptor. Constitutively expresses superfolder GFP under Pmax at the *Tn7* site. |
| BZB1011 *mrfp1*::Tn7 *ΔbtuB* | R[1] (Fig. 4) | This study | Resistant to group A colicins through deletion of the BtuB receptor. Constitutively expresses superfolder mRFP1 under Pmax at the *Tn7* site. |
| BZB1011 *mrfp1*::*Tn7* pC001-P*trc*:*immE2* | R[2] | This study | Resistant to colicin E2 by expression of ImmE2. Susceptible to all other colicins. |
| BZB1011 *mrfp1*::*Tn7* pTML9-P*max*:*immE2* | R[2] (Fig. 4) | This study | Resistant to colicin E2 by constitutive expression of ImmE2. Susceptible to all other colicins. |
| BZB1011 pColE2 | E2 (Fig 4) | [5] | Colicin E2-producing strain. Susceptible to all other colicins apart from E2. |
| BZB1011 *sfgfp*::*Tn7* pColE2 | E2 (Fig 3) | [1] | Colicin E2-producing strain. Susceptible to all other colicins apart from E2. Constitutively expresses superfolder GFP under P*max* at the Tn7 site. |

| | | | |
|---|---|---|---|
| BZB1011 *mrfp1*::*Tn7* pColE2 | E2 (Fig. 2) | [1] | Colicin E2-producing strain. Susceptible to all other colicins apart from E2. Constitutively expresses mRFP1 under P*max* at the Tn7 site. |
| BZB1011 *immE2*::*Tn7* pColE2 | E2* | [1] | Colicin E2-producing strain. Susceptible to all other colicins apart from E2. Constitutively expresses ImmE2 under P*max* at the Tn7 site. |
| BZB1011 *immE2*::*Tn7* pColE2 pBC11-P*ybaJ*:*ypet* | E2* | This study | Colicin E2-producing strain. Susceptible to all other colicins apart from E2. Constitutively expresses ImmE2 under P*max* at the Tn7 site. Constitutively expresses YPet. |
| BZB1011 *immE2*::*Tn7* pColE2 pBC43-P*ybaJ*:*blFP-Y3* | E2* (Fig. 3) | This study | Colicin E2-producing strain. Susceptible to all other colicins apart from E2. Constitutively expresses ImmE2 under P*max* at the Tn7 site. Constitutively expresses mNeonGreen. |
| BZB1011 *mrfp1*::*Tn7* pColE8 | E8 (Fig. 2,3) | [1] | Colicin E8-producing strain. Susceptible to all other colicins apart from E8. Constitutively expresses mRFP1 under P*max* at the Tn7 site. |
| BZB1011 *mrfp1*::*Tn7* pColE7 | E7 (Fig. 2,3) | [1] | Colicin E7-producing strain. Susceptible to all other colicins apart from E7. Constitutively expresses mRFP1 under P*max* at the Tn7 site. |
| BZB1011 *mrfp1*::*Tn7* pColA | A (Fig. 2,3) | [1] | Colicin A-producing strain. Susceptible to all other colicins apart from A Constitutively expresses mRFP1 under P*max* at the Tn7 site. |

**Extended Table 3.** Primers used in this study.

| Name | Sequence (5'-3') |
|---|---|
| TML-P5 | TTACAATGATTAAAAAAGCTGATTATGAGACAGTCTATGGCTACCAAACT |
| TML-P6 | CTGGAGCTCCACCGCGGTGGCGGCCGCTCTAGAACTAGTGGATCCGCCAGAATCCACCAGCCGGG |
| TML-P7 | CCCAGTCTCGAGGTCGACGGTATCGATAAGCTTGATATCGAATTCAACAACTTGTGGCGCAGAAC |
| TML-P8 | CCATAGACTGTCTCATAATCAGCTTTTTTAATCATTGTAAAGCATCCACAATAGA |
| TML-P9 | ATAGCAGGGAAACCACCGCC |
| TML-P10 | GCAGATTTTGCATCCGGGGC |
| TML-P57 | ACAGGGCTGAAATATGGGTATGCATTGCAGGCATGCAAG |
| TML-P58 | AGATCTCCGCAGCAGGAATTGGTGAAGACGAAAGGGCCTCGTGATACGC |
| TML-P59 | GAGGCCCTTTCGTCTTCACCAATTCCTGCTGCGGAGATCTG |
| TML-P60 | ATACTATGTTTCAGTTCCATCATTTTTTTTTTACCTCCTTAAACTCCTTATTTTGCATC |
| TML-P61 | AAGGAGTGAAAAAAAAAATGATGGAACTGAAACATAGTATTAGTGATTATACCGAG |
| TML-P62 | GCTTGCATGCCTGCAATGCATACCCATATTTCAGCCCTGTTTAAATCCT |
| oOPC-614 | GCTACCTGCTTTCTCTTTGCGCTTGC |
| oOPC-615 | TATGACCATGATTACGCCAAGCGCGC |

### Supplementary Methods

#### 1. Characterisation of the printing method

**1.1** Growth of *E. coli* within printed arrays

Once the printed arrays were in LB-Miller medium and incubated at 37 °C in a static incubator, the embedded *E. coli* grew from single-cell dispersions (with mean 2D centre-to-centre distances of 18 ± 2 μm) into 3D-microcolonies (Fig. 1e, f, and Extended Fig. 1a). After approximately 6 h of growth, *E. coli* invaded the surrounding medium (Extended Fig. 1e). Competition experiments were completed after 18 h because of nutrient depletion and saturation of the LB-Miller medium with *E. coli*. We considered that the cells in the medium surrounding the array might affect cells in the printed array via nutrient consumption or colicin production. However, we reasoned that any such effects from these cells, which are free to move and mix in the media, will only act to reduce the impacts of spatial structure in the array. We decided, therefore, to run our experiments with these cells present in the medium as it both made experiments simpler and represented a conservative test of the role of spatial structure in the array. And, as detailed in the main text, we did indeed see strong effects of spatial structure in the arrays, even though there were free cells in the media.

Compared to previous bacterial warfare experiments that spot colonies onto agar plates (5 μL), our printed communities are approximately 100 times smaller (by 2D area) (Fig. 1b, c). The communities were typically constructed from 7 x 8 x 1 droplets in the *x*, *y,* and *z* dimensions (1 layered structures) and swelled in medium to dimensions of 908 ± 65 x 998 ± 45 µm (*x*, *y*) (Extended Fig. 2a, b). We used 1-layered printed communities for competition experiments because of the simplicity of controlling genetic mixing by 2D patterns.

Within the printed arrays, after 18 h of growth, 3D microcolonies comprising of tightly packed chromosomally-labelled GFP and RFP BZB1011 strains had median diameters of 15 (10–21) µm, 2D aspect ratios (*x*, *y*) of 1.5 (1.3–1.8), and cross-sectional areas (*x*, *y*) of 156 (42–212) $\mu m^2$ (Extended Fig. 1b-d). These microcolonies excluded the agarose in the timeframe of experiments (Extended Fig. 1f).

1.2 Printing fidelity

For calculating cellular ratios (frequencies) within the printed arrays, estimates by flow cytometry, 2D segmentation, 3D segmentation, and colony-forming units (CFU), all gave the same results (Extended Fig. 2c). For simplicity, the frequencies of strains were calculated by 2D segmentation in most competition experiments.

With regards to the printing fidelity of patterned communities, the theoretical segregation index (*SI*) (see Methods) of printing maps matched well with the calculated *SI* of printed patterns. This correlation was consistent over different size regimes of the local neighbourhood. Overall, the *SI* values obtained from printing replicates were consistent with one another, which demonstrated the reproducibility of the technique (Fig. 1h). For competition experiments between two susceptible (S *gfp* and S *rfp*) strains in arrays printed at different *SI* values, but at the same starting density (1:1), 1:1 frequency ratios between strains were maintained at all *SI* values after 18 h of competition, and the morphologies of microcolonies were similar at different *SI* values (Fig. 1i, Extended Fig. 2d). This demonstrated that different printing patterns did not affect the affect the growth of similar genotypes that compete exploitatively (S *gfp* and S *rfp*).

#### 2. Colicins as a model for interference competition in bacteria

2.1 Colicin biology

*E. coli* can produce protein toxins that can target susceptible strains of *E. coli*. These toxins are called colicins and are critical for interference competition between different strains of *E. coli*. Genotypes that can produce colicins carry a small plasmid (pCol, type I plasmids: 6–10 kb) in about 20 copies per cell. Type I plasmids encode for group A colicins, which are classified as group A because they parasitise the Ton system in *E. coli* to enter the periplasm[6]. In this study, only group A colicins were used (colicin E2, E7, E8 and A). Colicin E2, E7, and E8 are nuclease colicins, which kill susceptible cells through DNA damage. Colicin A is a pore-forming colicin, which kills susceptible cells by disrupting the periplasm membrane potential. On the pCol plasmid, the operon that express the nuclease and pore-forming group A colicins is controlled by the protein LexA, which represses the SOS promoter for the operon[7]. Transcription of the colicin operon is controlled by the SOS response (a global repair response to DNA damage[8]). During the SOS response, the protein RecA is upregulated and activated by binding to single-stranded breaks. The activation of RecA enables self-cleavage of the LexA repressor allowing for binding of the RNA polymerase for transcription of the colicin operon (Extended Fig. 3a). Therefore, if a cell containing the pCol plasmid experiences DNA damage, it will express colicins.

For nuclease colicins in group A, the first gene encodes for the colicin itself, and the second gene downstream encodes for the cognate immunity protein (Extended Fig. 3b, c). The cognate immunity protein of nuclease colicins are controlled by the SOS promoter (LexA repressor) and is also constitutively expressed by a promoter sequence encoded in part of the colicin structure. This ensures the immunity protein is in excess so there is no self-intoxication by the nucleases. A susceptible cell to a colicin is a cell that does not express that colicin's cognate immunity protein. The last gene in the operon is the lysis gene, which encodes for the lysis protein required for release of the colicin. In pore-forming colicins, there is no gene encoding for the immunity protein in the operon. It is encoded on the opposite DNA strand under constitutive expression.

In summary, colicins are encoded on a small plasmid that contains an operon transcribing the colicin protein (toxin), immunity protein, and lysis protein required for release. When a colicin producer releases toxins, it sacrifices itself and dies.

#### 2.2 Strategy for generating colicin-resistant strains

Group A colicins enter the periplasm of susceptible *E. coli* cells by binding to the vitamin B12 outer-membrane receptor (BtuB) and being pulled through the outer membrane porin F (OmpF) by the Ton system (Fig 4b, Extended Fig. 4a). The binding to BtuB is critical because if *btuB* is deleted, the originally susceptible cells become resistant to any group A colicin (Fig 4b, Extended Fig. a, b). This is the common mechanism for resistance[9]. We created the resistant strain $R^1$ (BZB1011 *mrfp1::Tn7 ΔbtuB*) by deleting *btuB* (Fig. 4b, Extended Fig. 4a, b). Therefore, for $R^1$, there was no BtuB in the outer membrane.

Another mechanism of resistance to a specific group A colicin is through expression of the toxin's cognate immunity protein. Colicin-producers do not self-intoxicate because of the constitutive expression of a cognate immunity protein. Therefore, another mechanism to create resistance in a susceptible strain (without deleting *btuB*) is to introduce the constitutive expression of the toxin's cognate immunity protein. In this case, we created the strain $R^2$ (BZB1011 *mrfp1*::*Tn7* pC001-P*trc*:*immE2*, and BZB1011 *mrfp1*::*Tn7* pTML-9-P*max*:*immE2*), which carried plasmids that constitutively expressed the cognate immunity protein (ImmE2) for colicin E2. In this case, $R^2$ expresses BtuB and is resistant to colicin E2. For $R^2$, translocation of colicin E2 to the cytosol still occurs, however, colicin E2's DNase activity is neutralised by cytosolic immE2 (Fig 4b, Extended Fig. 4a, b). However, if other group A colicins translocate, the cytosolic ImmE2 would not confer resistance to the strain.

#### Supplementary Figures

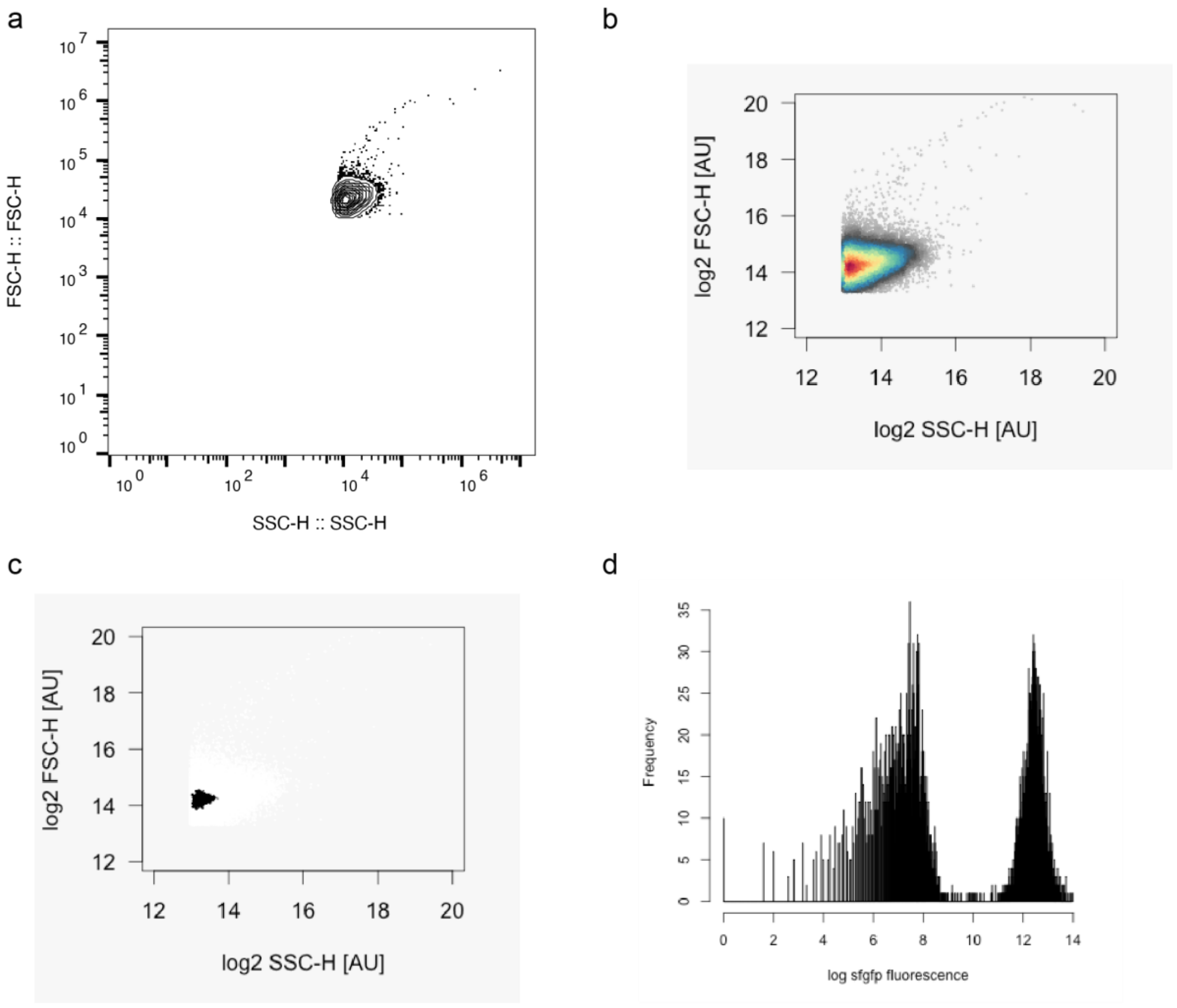


**Supplementary Fig. 1** Analysis pipeline of flow cytometry data in Extended Fig. 2c. An example of one biological replicate used to calculate the frequencies of BZB1011 *sfgfp*::*Tn7* and BZB1011 *mrfp1::Tn7* in printed arrays after 18 hours of competition. **a-b**, Contour plot (**a**) and dot plot (**b** – coloured by density) of the raw flow cytometry data with initial thresholds of 10 000 and 8000 for the forward and side scatters, respectively. **c**, A dot plot of the gated cell population based on the forward and side scatter. **d**, A histogram of the $\log_2$ fluorescence intensities of the cell population.

#### Supplementary Tables

**Supplementary Tables 1.** A statistical test for Fig. 1i: competition experiments between BZB1011 *sfgfp*::*Tn7* and BZB1011 *mrfp1*::*Tn7* in printed arrays. The tables show a statistical test (Kruskal-Wallis test) on the differences in median frequencies of BZB1011 *sfgfp*::*Tn7* in arrays printed at different *SI* values after 18 h of competition.

| Table Analysed | Fig 1i – Comparison of S *gfp* frequencies stats |
|---|---|
| Kruskal-Wallis test | |
| P value | 0.2736 |
| Exact or approximate P value? | Approximate |
| P value summary | ns |
| Do the medians vary signif. (P < 0.05)? | No |
| Number of groups | 5 |
| Kruskal-Wallis statistic | 5.137 |
| Data summary | |
| Number of treatments (columns) | 5 |
| Number of values (total) | 20 |

| | | | | | | |
|---|---|---|---|---|---|---|
| Number of families | 1 | | | | | |
| Number of comparisons per family | 10 | | | | | |
| Alpha | 0.05 | | | | | |
| Dunn's multiple comparisons test | Mean rank diff. | Significant? | Summary | Adjusted P Value | | |
| 0.94 vs. 0.88 | 2.583 | No | ns | >0.9999 | A-B | |
| 0.94 vs. 0.80 | -1.250 | No | ns | >0.9999 | A-C | |
| 0.94 vs. 0.67 | 6.250 | No | ns | >0.9999 | A-D | |
| 0.94 vs. 0.50 | 5.450 | No | ns | >0.9999 | A-E | |
| 0.88 vs. 0.80 | -3.833 | No | ns | >0.9999 | B-C | |
| 0.88 vs. 0.67 | 3.667 | No | ns | >0.9999 | B-D | |
| 0.88 vs. 0.50 | 2.867 | No | ns | >0.9999 | B-E | |
| 0.80 vs. 0.67 | 7.500 | No | ns | 0.7300 | C-D | |
| 0.80 vs. 0.50 | 6.700 | No | ns | 0.9136 | C-E | |
| 0.67 vs. 0.50 | -0.8000 | No | ns | >0.9999 | D-E | |
| Test details | Mean rank 1 | Mean rank 2 | Mean rank diff. | n1 | n2 | Z |
| 0.94 vs. 0.88 | 13.25 | 10.67 | 2.583 | 4 | 3 | 0.5717 |
| 0.94 vs. 0.80 | 13.25 | 14.50 | -1.250 | 4 | 4 | 0.2988 |
| 0.94 vs. 0.67 | 13.25 | 7.000 | 6.250 | 4 | 4 | 1.494 |
| 0.94 vs. 0.50 | 13.25 | 7.800 | 5.450 | 4 | 5 | 1.373 |
| 0.88 vs. 0.80 | 10.67 | 14.50 | -3.833 | 3 | 4 | 0.8484 |
| 0.88 vs. 0.67 | 10.67 | 7.000 | 3.667 | 3 | 4 | 0.8115 |
| 0.88 vs. 0.50 | 10.67 | 7.800 | 2.867 | 3 | 5 | 0.6635 |
| 0.80 vs. 0.67 | 14.50 | 7.000 | 7.500 | 4 | 4 | 1.793 |
| 0.80 vs. 0.50 | 14.50 | 7.800 | 6.700 | 4 | 5 | 1.688 |
| 0.67 vs. 0.50 | 7.000 | 7.800 | -0.8000 | 4 | 5 | 0.2016 |

**Supplementary Table 2.** A statistical test for Fig. 2c competition experiments between BZB1011 *sfgfp*::*Tn7* and BZB1011 *mrfp1*::*Tn7* pColE7 in printed arrays. The table shows a statistical test (Kruskal-Wallis test) on the differences in median frequencies of BZB1011 *sfgfp*::*Tn7* in arrays printed at different *SI* values after 18 h of competition.

| Table Analysed | Fig. 2a |
|---|---|
| Kruskal-Wallis test | |
| P value | <0.0001 |
| Exact or approximate P value? | Exact |
| P value summary | **** |
| Do the medians vary signif. (P < 0.05)? | Yes |
| Number of groups | 4 |
| Kruskal-Wallis statistic | 15.63 |
| Data summary | |
| Number of treatments (columns) | 4 |
| Number of values (total) | 18 |

**Supplementary Tables 3.** Statistical tests for Fig. 2e–h: competition experiments between BZB1011 *sfgfp*::*Tn7* and BZB1011 *mrfp1*::*Tn7* pColA (**e**), BZB1011 *sfgfp*::*Tn7* and BZB1011 *mrfp1*::*Tn7* pColE2 (**f**), BZB1011 *sfgfp*::*Tn7* and BZB1011 *mrfp1*::*Tn7* pColE8 (**g**), and BZB1011 *mrfp1*::*Tn7* pColE2 and BZB1011 *sfgfp*::*Tn7 ΔbtuB* in printed arrays (**f**). The tables show statistical tests on the changes in the median frequencies between competing strains at different *SI* values (unpaired Mann Whitney tests) and on the differences in frequencies between competing strains at the same *SI* value (Wilcoxon tests), after 18 h of competition.

| Table Analysed | Fig. 2 |
|---|---|
| Column B | Fig. 2e S vs A (SI = 0.94) |
| vs | vs |
| Column A | Fig. 2e S vs A (SI = 0.50) |
| Mann Whitney test | |
| P value | 0.6286 |
| Exact or approximate P value? | Exact |
| P value summary | ns |
| Significantly different (P < 0.05)? | No |
| One- or two-tailed P value? | Two-tailed |
| Sum of ranks in column A,B | 10, 18 |
| Mann-Whitney U | 4 |
| Difference between medians | |
| Median of column A | -0.2751, *n* = 3 |
| Median of column B | -0.08756, *n* = 4 |
| Difference: Actual | 0.1875 |
| Difference: Hodges-Lehmann | 0.1875 |

| Table Analysed | Fig. 2 |
|---|---|
| Column D | Fig. 2f S vs E2 (SI = 0.94) |
| vs | vs |
| Column C | Fig. 2f S vs E2 (SI = 0.50) |
| Mann Whitney test | |

| | |
|---|---|
| P value | 0.0020 |
| Exact or approximate P value? | Exact |
| P value summary | ** |
| Significantly different ($P < 0.05$)? | Yes |
| One- or two-tailed P value? | Two-tailed |
| Sum of ranks in column C,D | 10, 95 |
| Mann-Whitney U | 0 |
| Difference between medians | |
| Median of column C | -0.9996, $n = 4$ |
| Median of column D | -0.4110, $n = 10$ |
| Difference: Actual | 0.5886 |
| Difference: Hodges-Lehmann | 0.5887 |

| Table Analysed | Fig. 2 |
|---|---|
| Column F | Fig. 2g S vs E8 (SI = 0.94) |
| vs | vs |
| Column E | Fig. 2g S vs E8 (SI = 0.50) |
| Mann Whitney test | |
| P value | 0.4462 |
| Exact or approximate P value? | Exact |
| P value summary | ns |
| Significantly different ($P < 0.05$)? | No |
| One- or two-tailed P value? | Two-tailed |
| Sum of ranks in column E,F | 60.50, 75.50 |
| Mann-Whitney U | 24.50 |
| Difference between medians | |
| Median of column E | -1.000, $n = 8$ |
| Median of column F | -1.000, $n = 8$ |
| Difference: Actual | 0.000 |
| Difference: Hodges-Lehmann | 0.000 |

| Table Analysed | Fig. 2 |
|---|---|
| Column H | Fig. 2g E2 vs R2 (SI = 0.94) |
| vs | vs |
| Column G | Fig. 2g E2 vs R2 (SI = 0.50) |
| Mann Whitney test | |
| P value | 0.0336 |
| Exact or approximate P value? | Exact |
| P value summary | * |
| Significantly different ($P < 0.05$)? | Yes |
| One- or two-tailed P value? | Two-tailed |
| Sum of ranks in column G,H | 49, 42 |
| Mann-Whitney U | 4 |
| Difference between medians | |
| Median of column G | -0.08480, $n = 9$ |
| Median of column H | 0.06940, $n = 4$ |
| Difference: Actual | 0.1542 |

| | |
|---|---|
| Difference: Hodges-Lehmann | 0.1472 |

| One sample t and Wilcoxon test | Fig. 2g E2 vs R2 (SI = 0.50) |
|---|---|
| Theoretical median | 0.000 |
| Actual median | -0.08480 |
| Number of values | 9 |
| Wilcoxon Signed Rank Test | |
| Sum of signed ranks (W) | -45.00 |
| Sum of positive ranks | 0.000 |
| Sum of negative ranks | -45.00 |
| P value (two tailed) | 0.0039 |
| Exact or estimate? | Exact |
| P value summary | ** |
| Significant (alpha = 0.05)? | Yes |
| How big is the discrepancy? | |
| Discrepancy | -0.08480 |
| 95% confidence interval | -0.1500 to -0.05180 |
| Actual confidence level | 96.09 |

| One sample t and Wilcoxon Wilcoxon test | Fig. 2g E2 vs R2 (SI = 0.94) |
|---|---|
| Theoretical median | 0.000 |
| Actual median | 0.06940 |
| Number of values | 4 |
| Wilcoxon Signed Rank Test | |
| Sum of signed ranks (W) | 4.000 |
| Sum of positive ranks | 7.000 |
| Sum of negative ranks | -3.000 |
| P value (two tailed) | 0.6250 |
| Exact or estimate? | Exact |
| P value summary | ns |
| Significant (alpha = 0.05)? | No |
| How big is the discrepancy? | |
| Discrepancy | 0.06940 |
| 95% confidence interval | -0.07900 to 0.09360 |
| Actual confidence level | 87.50 |

**Supplementary Tables 4.** Statistical tests for Fig. 3: competition experiments between BZB1011 *sfgfp*::*Tn7* pColE2 and BZB1011 *mrfp1*::*Tn7* pColE7 (**a**), BZB1011 *sfgfp*::*Tn7* pColE2 and BZB1011 *mrfp1*::*Tn7* pColA (**b**), BZB1011 *sfgfp*::*Tn7* pColE2 and BZB1011 *mrfp1*::*Tn7* pColE8 (**c**), and BZB1011 *immE2*::*Tn7* pColE2 pBC43-P*ybaJ*:*blFP-Y3* and BZB1011 *mrfp1*::*Tn7* pColE8 in printed arrays (**d**). The tables show statistical tests on the changes in the median frequencies between competing strains at different *SI* values (unpaired Mann-Whitney tests) after 18 h of competition.

| Table Analysed | Fig. 3 |
|---|---|
| Column B | Fig, 3a E2 vs E7 (SI = 0.94) |
| vs | vs |

| | |
|---|---|
| Column A | Fig. 3a E2 vs E7 (SI = 0.50) |
| Mann Whitney test | |
| P value | 0.0286 |
| Exact or approximate P value? | Exact |
| P value summary | * |
| Significantly different (P < 0.05)? | Yes |
| One- or two-tailed P value? | Two-tailed |
| Sum of ranks in column A,B | 22, 6 |
| Mann-Whitney U | 0 |
| Difference between medians | |
| Median of column A | 1.000, *n* = 4 |
| Median of column B | 0.6819, *n* = 3 |
| Difference: Actual | -0.3181 |
| Difference: Hodges-Lehmann | -0.3181 |

| | |
|---|---|
| Table Analysed | Fig. 3 |
| Column D | Fig. 3b E2 vs A (SI = 0.94) |
| vs | vs |
| Column C | Fig. 3b E2 vs A (SI = 0.50) |
| Mann Whitney test | |
| P value | >0.9999 |
| Exact or approximate P value? | Exact |
| P value summary | ns |
| Significantly different (P < 0.05)? | No |
| One- or two-tailed P value? | Two-tailed |
| Sum of ranks in column C,D | 42, 49 |
| Mann-Whitney U | 21 |
| Difference between medians | |
| Median of column C | -1.000, *n* = 6 |
| Median of column D | -1.000, *n* = 7 |
| Difference: Actual | 0.000 |
| Difference: Hodges-Lehmann | 0.000 |

| | |
|---|---|
| Table Analysed | Fig. 3 |
| Column F | Fig. 3c E2 vs E8 (0.94) |
| vs | vs |
| Column E | Fig. 3c E2 vs E8 (SI = 0.50) |
| Mann Whitney test | |
| P value | 0.0286 |
| Exact or approximate P value? | Exact |
| P value summary | * |
| Significantly different (P < 0.05)? | Yes |
| One- or two-tailed P value? | Two-tailed |
| Sum of ranks in column E,F | 10, 18 |
| Mann-Whitney U | 0 |
| Difference between medians | |
| Median of column E | -1.000, *n* = 4 |

| | |
|---|---|
| Median of column F | -0.06570, *n* = 3 |
| Difference: Actual | 0.9343 |
| Difference: Hodges-Lehmann | 0.9343 |

| | |
|---|---|
| Table Analysed | Fig. 3 |
| Column H | Fig. 3d E2* vs E8 (SI = 0.94) |
| vs | vs |
| Column G | Fig. 3d E2* vs E8* (SI = 0.50) |
| Mann Whitney test | |
| P value | 0.0337 |
| Exact or approximate P value? | Exact |
| P value summary | * |
| Significantly different (P < 0.05)? | Yes |
| One- or two-tailed P value? | Two-tailed |
| Sum of ranks in column G,H | 94 , 42 |
| Mann-Whitney U | 14 |
| Difference between medians | |
| Median of column G | 0.000, *n* = 9 |
| Median of column H | -1.000, *n* = 7 |
| Difference: Actual | -1.000 |
| Difference: Hodges-Lehmann | -1.000 |

**Supplementary Table 5.** A statistical test for Fig. 4c: competition experiments between BZB1011 *sfgfp*::*Tn7* and BZB1011 *mrfp1*::*Tn7* and BZB1011 in structured printed arrays. The tables show statistical tests (Wilcoxon tests) on the difference in the median productivities of BZB1011 *sfgfp*::*Tn7* and BZB1011 *mrfp1*::*Tn7* after 18 h of competition.

| | |
|---|---|
| Table Analysed (Wilcoxon test) | Fig. 4c |
| Column D | S-blue |
| vs | vs |
| Column C | S-orange |
| Wilcoxon matched-pairs signed rank test | |
| P value | >0.9999 |
| Exact or approximate P value? | Exact |
| P value summary | ns |
| Significantly different (P < 0.05)? | No |
| One- or two-tailed P value? | Two-tailed |
| Sum of positive, negative ranks | 14.00, -14.00 |
| Sum of signed ranks (W) | 0.000 |
| Number of pairs | 7 |
| Number of ties (ignored) | 0 |
| Median of differences | |
| Median | -5.400 |
| How effective was the pairing? | |
| rs (Spearman) | -0.03571 |
| P value (one tailed) | 0.4817 |
| P value summary | ns |
| Was the pairing significantly effective? | No |

**Supplementary Table 6.** A statistical test for Fig. 4d: competition experiments between BZB1011 *sfgfp*::*Tn7* and BZB1011 *mrfp1*::*Tn7* and BZB1011 pColE2 in structured printed arrays. The table shows a statistical test (Wilcoxon test) on the difference in the median productivities of BZB1011 *sfgfp*::*Tn7* and BZB1011 *mrfp1*::*Tn7* after 18 h of competition.

| Table Analysed (Wilcoxon test) | Fig. 4d |
|---|---|
| Column F | S-blue |
| vs | vs |
| Column E | S-orange |
| Wilcoxon matched pairs signed rank test | |
| P value | 0.0002 |
| Exact or approximate P value? | Exact |
| P value summary | *** |
| Significantly different (P < 0.05)? | Yes |
| One- or two-tailed P value? | Two-tailed |
| Sum of positive, negative ranks | 0.000, -91.00 |
| Sum of signed ranks (W) | -91.00 |
| Number of pairs | 13 |
| Number of ties (ignored) | 0 |
| Median of differences | |
| Median | -50.80 |
| How effective was the pairing? | |
| rs (Spearman) | 0.5386 |
| P value (one-tailed) | 0.0299 |
| P value summary | * |
| Was the pairing significantly effective? | Yes |

**Supplementary Table 7.** Statistical tests for Fig. 4e and d, and Fig. 4e and f: competition experiments between BZB1011 *sfgfp*::*Tn7* and BZB1011 pColE2 (Fig. 4e); BZB1011 *sfgfp*::*Tn7* and BZB1011 *mrfp1*::*Tn7* and BZB1011 pColE2 (Fig. 4d), and BZB1011 *sfgfp*::*Tn7* and BZB1011 *mrfp1*::*Tn7 ΔbtuB* and BZB1011 pColE2 (Fig. 4h) in structured printed arrays. The table shows statistical tests (Mann-Whitney test) on the difference in median productivities of BZB1011 *sfgfp*::*Tn7* in Fig. 4e and d, and Fig. 4e and f after 18 h of competition.

| Table Analysed (Mann-Whitney test) | Comparison of S |
|---|---|
| Column C | S-orange (Fig. 4e) |
| vs. | vs. |
| Column B | S-orange (Fig. 4d) |
| Mann Whitney test | |
| P value | 0.0029 |
| Exact or approximate P value? | Exact |
| P value summary | ** |
| Significantly different (P < 0.05)? | Yes |
| One- or two-tailed P value? | Two-tailed |
| Sum of ranks in column B,C | 141 , 12 |
| Mann-Whitney U | 2 |
| Difference between medians | |

| | |
|---|---|
| Median of column B | 0.5927, n = 13 |
| Median of column C | 0.01523, n = 4 |
| Difference: Actual | -0.5774 |
| Difference: Hodges-Lehmann | -0.5529 |

| Table Analysed (Mann-Whitney test) | Fig. 4 |
|---|---|
| Column D | S-orange (Fig. 4f) |
| vs | vs |
| Column C | S-orange (Fig. 4e) |
| Mann Whitney test | |
| P value | 0.1143 |
| Exact or approximate P value? | Exact |
| P value summary | ns |
| Significantly different (P < 0.05)? | No |
| One- or two-tailed P value? | Two-tailed |
| Sum of ranks in column C,D | 24, 12 |
| Mann-Whitney U | 2 |
| Difference between medians | |
| Median of column C | 1.523, *n* = 4 |
| Median of column D | 0.6637, *n* = 4 |
| Difference: Actual | -0.8589 |
| Difference: Hodges-Lehmann | -0.6723 |

**Supplementary Table 8.** A statistical test for Fig. 4d and f: competition experiments between BZB1011 *sfgfp*::*Tn7* and BZB1011 *mrfp1*::*Tn7* and BZB1011 pColE2 (Fig. 4d); and BZB1011 *sfgfp*::*Tn7* and BZB1011 *mrfp1*::*Tn7 ΔbtuB* and BZB1011 pColE2 (Fig. 4h) in structured printed arrays. Table shows a statistical test (Mann-Whitney test) on the difference in median productivities of BZB1011 *sfgfp*::*Tn7* in Fig. 4d and 4f after 18 h of competition.

| Table Analysed (Mann-Whitney test) | Fig. 4 |
|---|---|
| Column G | S-orange (Fig. 4d) |
| vs | vs |
| Column E | S-orange (Fig. 4f) |
| Mann Whitney test | |
| P value | 0.0008 |
| Exact or approximate P value? | Exact |
| P value summary | *** |
| Significantly different (P < 0.05)? | Yes |
| One- or two-tailed P value? | Two-tailed |
| Sum of ranks in column E, G | 143, 10 |
| Mann-Whitney U | 0 |
| Difference between medians | |
| Median of column E | 59.27, *n* = 13 |
| Median of column G | 0.6637, *n* = 4 |
| Difference: Actual | -58.60 |
| Difference: Hodges-Lehmann | -58.60 |

**Supplementary Table 9.** A statistical test for Fig. 4d and g: competition experiments between BZB1011 *sfgfp*::*Tn7* and BZB1011 *mrfp1*::*Tn7* and BZB1011 pColE2 (Fig. 4d); and BZB1011 *sfgfp*::*Tn7* and BZB1011 *mrfp1*::*Tn7 immE2* and BZB1011 pColE2 (Fig. 4g) in structured printed arrays. The table shows a statistical test (Mann-Whitney test) on the difference in median productivities of BZB1011 *sfgfp*::*Tn7* in Fig. 4d and 4g after 18 h of competition.

| Table Analysed (Mann-Whitney test) | Fig. 4 |
|---|---|
| Column E | S-orange (Fig. 4f) |
| vs. | vs. |
| Column B | S-orange (Fig. 4d) |
| Mann Whitney test | |
| P value | 0.6404 |
| Exact or approximate P value? | Exact |
| P value summary | ns |
| Significantly different (P < 0.05)? | No |
| One- or two-tailed P value? | Two-tailed |
| Sum of ranks in column B, E | 185, 166 |
| Mann-Whitney U | 75 |
| Difference between medians | |
| Median of column B | 59.27, n = 13 |
| Median of column E | 47.16, n = 13 |
| Difference: Actual | -12.11 |
| Difference: Hodges-Lehmann | -7.630 |

**Supplementary Tables 10.** Statistical tests for Extended Fig. 2b: measurements of the *x* and *y* dimensions in printed arrays formed from droplet networks with dimensions 7 x 8 x 1 and 8 x 8 x 1 in the *x*, *y*, and *z* dimensions. Includes tests for normal distributions of the *x* and *y* dimensions and statistical tests (Welch's t tests) on the differences in the mean *x* and *y* dimensions in 7 x 8 x 1 and 8 x 8 x 1 printed arrays.

| Test for normal distribution | | | | |
|---|---|---|---|---|
| Anderson-Darling test | | | | |
| A2* | 0.1988 | 0.5347 | 0.2450 | 0.3232 |
| P value | 0.8503 | 0.1343 | 0.6977 | 0.4767 |
| Passed normality test (alpha=0.05)? | Yes | Yes | Yes | Yes |
| P value summary | ns | ns | ns | ns |
| D'Agostino & Pearson test | | | | |
| K2 | 0.8846 | 1.850 | 0.7671 | 0.2518 |
| P value | 0.6425 | 0.3964 | 0.6814 | 0.8817 |
| Passed normality test (alpha=0.05)? | Yes | Yes | Yes | Yes |
| P value summary | ns | ns | ns | ns |
| Shapiro-Wilk test | | | | |
| W | 0.9608 | 0.8928 | 0.9532 | 0.9500 |
| P value | 0.7951 | 0.1281 | 0.6837 | 0.6372 |
| Passed normality test (alpha=0.05)? | Yes | Yes | Yes | Yes |
| P value summary | ns | ns | ns | ns |
| Kolmogorov-Smirnov test | | | | |
| KS distance | 0.1459 | 0.2182 | 0.1316 | 0.1573 |

| | | | | |
|---|---|---|---|---|
| P value | > 0.1000 | > 0.1000 | > 0.1000 | > 0.1000 |
| Passed normality test (alpha=0.05)? | Yes | Yes | Yes | Yes |
| P value summary | ns | ns | ns | ns |
| Number of values | 12 | 12 | 12 | 12 |

| Table Analysed (Welch’s t test) | 7 x 8 x 1 |
|---|---|
| Column D | y 7 |
| vs. | vs. |
| Column C | x 7 |
| Unpaired t test with Welch's correction | |
| P value | 0.0010 |
| P value summary | *** |
| Significantly different (P < 0.05)? | Yes |
| One- or two-tailed P value? | Two-tailed |
| Welch-corrected t, df | t = 3.871, df = 19.55 |
| How big is the difference? | |
| Mean of column C | 908.8 |
| Mean of column D | 997.7 |
| Difference between means (D - C) ± SEM | 88.91 ± 22.97 |
| 95% confidence interval | 40.92 to 136.9 |
| R squared (eta squared) | 0.4338 |
| F test to compare variances | |
| F, DFn, Dfd | 2.094, 11, 11 |
| P value | 0.2359 |
| P value summary | ns |
| Significantly different (P < 0.05)? | No |
| Data analysed | |
| Sample size, column C | 12 |
| Sample size, column D | 12 |

| Table Analysed (Welch’s t test) | 8 x 8 x 1 |
|---|---|
| Column B | y 8 |
| vs | vs. |
| Column A | x 8 |
| Unpaired t test with Welch's correction | |
| P value | 0.6072 |
| P value summary | ns |
| Significantly different (P < 0.05)? | No |
| One- or two-tailed P value? | Two-tailed |
| Welch-corrected t, df | t=0.5223, df=20.05 |
| How big is the difference? | |
| Mean of column A | 936.3 |
| Mean of column B | 952.3 |
| Difference between means (B - A) ± SEM | 16.04 ± 30.71 |
| 95% confidence interval | -48.01 to 80.08 |
| R squared (eta squared) | 0.01342 |
| F test to compare variances | |
| F, DFn, Dfd | 1.905, 11, 11 |
| P value | 0.3001 |

| | |
|---|---|
| P value summary | ns |
| Significantly different (P < 0.05)? | No |
| Data analysed | |
| Sample size, column A | 12 |
| Sample size, column B | 12 |

**Supplementary Table 11.** A statistical test for Extended Fig. 2c: competition experiments between BZB1011 *sfgfp*::*Tn7* and BZB1011 *mrfp1*::*Tn7* in arrays printed at an *SI* = 0.50 with different measurements used to calculate the strain frequencies after 18 h of competition. The table shows a statistical test (Kruskal-Wallis) on the differences in the median frequencies of BZB1011 *sfgfp*::*Tn7* calculated by different measurements (2D and 3D segmentation, flow cytometry and CFU).

| Table Analysed | Extended Fig. 2c |
|---|---|
| Kruskal-Wallis test | |
| P value | 0.2415 |
| Exact or approximate P value? | Exact |
| P value summary | ns |
| Do the medians vary signif. (P < 0.05)? | No |
| Number of groups | 4 |
| Kruskal-Wallis statistic | 4.306 |
| Data summary | |
| Number of treatments (columns) | 4 |
| Number of values (total) | 18 |

**Supplementary Table 12.** A statistical test for Extended Fig. 2d: competition experiments between BZB1011 *sfgfp*::*Tn7* and BZB1011 *mrfp1*::*Tn7* in printed arrays. The table shows a statistical test (Kruskal-Wallis) on the differences in median lengths (diameter of *z*-projected microcolonies) of microcolonies of BZB1011 *sfgfp*::*Tn7* in arrays printed at different *SI* values after 18 h of competition.

| Table Analysed | length |
|---|---|
| Kruskal-Wallis test | |
| P value | 0.4247 |
| Exact or approximate P value? | Approximate |
| P value summary | ns |
| Do the medians vary signif. (P < 0.05)? | No |
| Number of groups | 5 |
| Kruskal-Wallis statistic | 3.864 |
| Data summary | |
| Number of treatments (columns) | 5 |
| Number of values (total) | 20 |

**Supplementary Tables 13.** Statistical tests for Extended Fig. 3i: competition experiments between BZB1011 *sfgfp*::*Tn7* and BZB1011 *mrfp1*::*Tn7* pColE7 in printed arrays. The tables show statistical tests (Mann Whitney tests) on the differences in the median frequencies of BZB1011 *sfgfp*::*Tn7* measured by 2D segmentation (from *z*-projected confocal images) and CFU in arrays printed at *SI* = 0.50 and 0.94 after 18 h of competition.

| Table Analysed | *SI* = 0.50 |
|---|---|
| Column B | CFU |
| vs | vs. |
| Column A | 2D |
| Mann Whitney test | |
| P value | 0.7308 |
| Exact or approximate P value? | Exact |
| P value summary | ns |
| Significantly different (P < 0.05)? | No |
| One- or two-tailed P value? | Two-tailed |
| Sum of ranks in column A, B | 39, 52 |
| Mann-Whitney U | 18 |
| Difference between medians | |
| Median of column A | 23.43, n = 6 |
| Median of column B | 27.12, n = 7 |
| Difference: Actual | 3.694 |
| Difference: Hodges-Lehmann | 2.249 |

| Table Analysed | *SI* = 0.94 |
|---|---|
| Column D | CFU |
| vs | vs. |
| Column C | 2D |
| Mann Whitney test | |
| P value | 0.5887 |
| Exact or approximate P value? | Exact |
| P value summary | ns |
| Significantly different (P < 0.05)? | No |
| One- or two-tailed P value? | Two-tailed |
| Sum of ranks in column C,D | 43, 35 |
| Mann-Whitney U | 14 |
| Difference between medians | |
| Median of column C | 61.00, n = 6 |
| Median of column D | 60.25, n = 6 |
| Difference: Actual | -0.7500 |
| Difference: Hodges-Lehmann | -0.7850 |

**Supplementary Table 14.** Statistical tests for Extended Fig. 4c: competition experiments between BZB1011 *sfgfp*::*Tn7* and BZB1011 *mrfp1*::*Tn7 ΔbtuB* in printed arrays. The tables show statistical tests on the changes in the median frequencies between competing strains at different *SI* values (unpaired Mann-Whitney tests) and on the differences in frequencies between competing strains at the same *SI* value (Wilcoxon tests), after 18 h of competition.

| Table Analysed | Fig. 2 and Extended Fig. 3c |
|---|---|
| Column J | Ext Fig. 3c S vs R1 (SI = 0.94) |
| vs | vs. |
| Column I | Ext Fig. 3c S vs R1 (SI = 0.50) |
| Mann Whitney test | |
| P value | 0.0043 |
| Exact or approximate P value? | Exact |

| | |
|---|---|
| P value summary | ** |
| Significantly different (P < 0.05)? | Yes |
| One- or two-tailed P value? | Two-tailed |
| Sum of ranks in column I,J | 56.50 , 21.50 |
| Mann-Whitney U | 0.5000 |
| Difference between medians | |
| Median of column I | 0.1448, n = 6 |
| Median of column J | 0.01320, n = 6 |
| Difference: Actual | -0.1316 |
| Difference: Hodges-Lehmann | -0.1652 |

| One sample t and Wilcoxon test | Ext Fig. 3c S vs R1 (SI = 0.50) |
|---|---|
| Theoretical median | 0.000 |
| Actual median | 0.1448 |
| Number of values | 6 |
| Wilcoxon Signed Rank Test | |
| Sum of signed ranks (W) | 21.00 |
| Sum of positive ranks | 21.00 |
| Sum of negative ranks | 0.000 |
| P value (two tailed) | 0.0312 |
| Exact or estimate? | Exact |
| P value summary | * |
| Significant (alpha=0.05)? | Yes |
| How big is the discrepancy? | |
| Discrepancy | 0.1448 |
| 95% confidence interval | 0.1080 to 0.3532 |
| Actual confidence level | 96.88 |

| Once sample t and Wilcoxon test | Ext Fig. 3c S vs R1 (SI = 0.94) |
|---|---|
| Theoretical median | 0.000 |
| Actual median | 0.01320 |
| Number of values | 6 |
| Wilcoxon Signed Rank Test | |
| Sum of signed ranks (W) | 3.000 |
| Sum of positive ranks | 12.00 |
| Sum of negative ranks | -9.000 |
| P value (two tailed) | 0.8438 |
| Exact or estimate? | Exact |
| P value summary | ns |
| Significant (alpha=0.05)? | No |
| How big is the discrepancy? | |
| Discrepancy | 0.01320 |
| 95% confidence interval | -0.1206 to 0.1080 |
| Actual confidence level | 96.88 |
